# Interactions with presynaptic photoreceptors mediated by the Dpr11 and DIP-γ cell surface proteins control selection and survival of *Drosophila* amacrine neurons

**DOI:** 10.1101/679704

**Authors:** Kaushiki P. Menon, Vivek Kulkarni, Shin-ya Takemura, Michael Anaya, Kai Zinn

**Affiliations:** Division of Biology and Biological Engineering, California Institute of Technology, Pasadena, CA 91125; Janelia Research Campus, Howard Hughes Medical Institute, Ashburn, VA 20147

## Abstract

*Drosophila* R7 UV photoreceptors (PRs) are divided into yellow (y) and pale (p) subtypes with different wavelength sensitivities. yR7 PRs express the Dpr11 cell surface protein and are presynaptic to Dm8 amacrine neurons (yDm8) that express Dpr11’s binding partner DIP-γ, while pR7 PRs synapse onto DIP-γ-negative pDm8 neurons. Dpr11 and DIP-γ expression patterns define yellow and pale medulla color vision circuits that project to higher-order areas. *DIP-* γ and *dpr11* mutations affect the morphology of yDm8 arbors in the yellow circuit. yDm8 neurons are generated in excess during development and compete for presynaptic yR7 partners. Transsynaptic interactions between Dpr11 and DIP-γ are required for generation of neurotrophic signals that allow yDm8 neurons to survive. yDm8 and pDm8 neurons do not normally compete for neurotrophic support, but can be forced to do so by manipulating R7 subtype fates. DIP-γ-Dpr11 interactions allow yDm8 neurons to select yR7 PRs as their home column partners.

## INTRODUCTION

The chemoaffinity hypothesis (Sperry, 1963) proposed that assembly of neural circuits involves interactions among cell-surface proteins (CSPs) that label individual neurons or neuronal types. This hypothesis applies well to systems such as the *Drosophila* brain and vertebrate retina in which synaptic connectivity is largely controlled by intrinsic gene expression patterns. There are approximately 1000 CSPs encoded in the *Drosophila* genome that are likely to be involved in cell-cell recognition events (Kurusu et al., 2008). In the pupal visual system, (Tan et al., 2015) characterized CSP gene expression in R7 and R8 photoreceptors (PRs) and the five types of lamina neurons (L1-L5), which relay information from R1-R6 PRs. Each of these seven neuronal types expresses more than 250 of the 1000 CSP genes, and each type differs from any of the others by expression of least 50 CSP genes.

To find the CSPs in these cellular expression profiles that are likely to be important for circuit assembly, we and others have focused on interaction partners expressed on synaptically connected neurons. The “Dpr-ome” is a network of immunoglobulin superfamily (IgSF) CSPs that was discovered in an *in vitro* “interactome” screen of all *Drosophila* IgSF proteins(Carrillo et al., 2015; Özkan et al., 2013). The current Dpr-ome has 21 2-IgSF Dpr proteins (Nakamura et al., 2002), each of which binds to one or more of the 11 3-IgSF DIP proteins. Most DIPs interact with multiple Dprs and *vice versa*, and their binding affinities vary between 1 and 200 µM (Cosmanescu et al., 2018). Each DIP and Dpr is expressed in a unique subset of neurons at each stage of development. In the visual system, neurons expressing a DIP tend to be postsynaptic to neurons that express a Dpr to which that DIP binds *in vitro* (Carrillo et al., 2015; Cosmanescu et al., 2018; Davis et al., 2018; Tan et al., 2015; Xu et al., 2018b). DIPs and Dprs define an IgSF family, present in both protostomes and deuterostomes, that has been denoted as the Wirins. The five mammalian IgLON proteins are members of the Wirin family (Cheng et al., 2019b).

Genetic analysis of DIP-Dpr pairs has revealed a variety of phenotypes. DIP-γ and Dpr11 regulate neuromuscular junction (NMJ) morphology in larvae, and control survival of DIP-γ-expressing postsynaptic cells in the pupal optic lobe (Carrillo et al., 2015; Xu et al., 2018b). Interactions between postsynaptic DIP-α and presynaptic Dprs 6 and 10 also control survival of postsynaptic optic lobe neurons, and can determine their synaptic connectivity patterns (Xu et al., 2018b). In the larval and adult neuromuscular systems, however, DIP-α and Dpr10 control branching of DIP-α-expressing motor axons onto muscle fibers that express Dpr10 (Ashley et al., 2019; Cheng et al., 2019a; Venkatasubramanian et al., 2019). Dprs and DIPs regulate fasciculation and sorting of olfactory receptor neuron axons in the antennal lobe (Barish et al., 2018). In the lamina of the optic lobe, DIPs prevent ectopic synapse formation (Xu et al., 2018a).

DIPs and Dprs may be components of large CSP repertoires that confer a unique surface identity to each type of neuron. Each neuron uses its repertoire to sculpt its morphology, determine its synaptic connectivity, and regulate its physiological properties. The total number of CSP genes is only about fourfold larger than the size of a repertoire (Tan et al., 2015), so the repertoires of different neurons necessarily overlap. Two neuronal types might have some of the same CSPs in their repertoires but use them in different ways, depending on their developmental needs.

Here we show that Dpr11 and DIP-γ expression patterns define a labeled-line color vision circuit that includes R7 PRs and Dm8 amacrine neurons in the medulla of the optic lobe. R7 and R8 transmit chromatic information from the retina to the medulla. R7 is a UV receptor, while R8 responds to visible light. Each ommatidium in the compound eye contains an R7, an R8, and six achromatic PRs (R1-R6) that transmit information relevant to motion detection (Figure 1A). R1-R6 synapse onto lamina neurons, which in turn project to layers M1-M5 of the medulla. R7 and R8 axons grow through the lamina and into the medulla, where they terminate in layers M6 (R7) and M3 (R8) (Figure 1C). The medulla is a ten-layered neuropil that is divided into columns, each of which roughly corresponds to one of the ∼750 ommatidia of the compound eye. It contains about 100 types of neurons. Some of these arborize only in the medulla, either in single columns or across multiple columns, while others have dendrites in the medulla and project to higher-order visual areas, including the lobula and lobula plate (reviewed by (Hadjieconomou et al., 2011; Sanes and Zipursky, 2010)).

**Figure 1:**
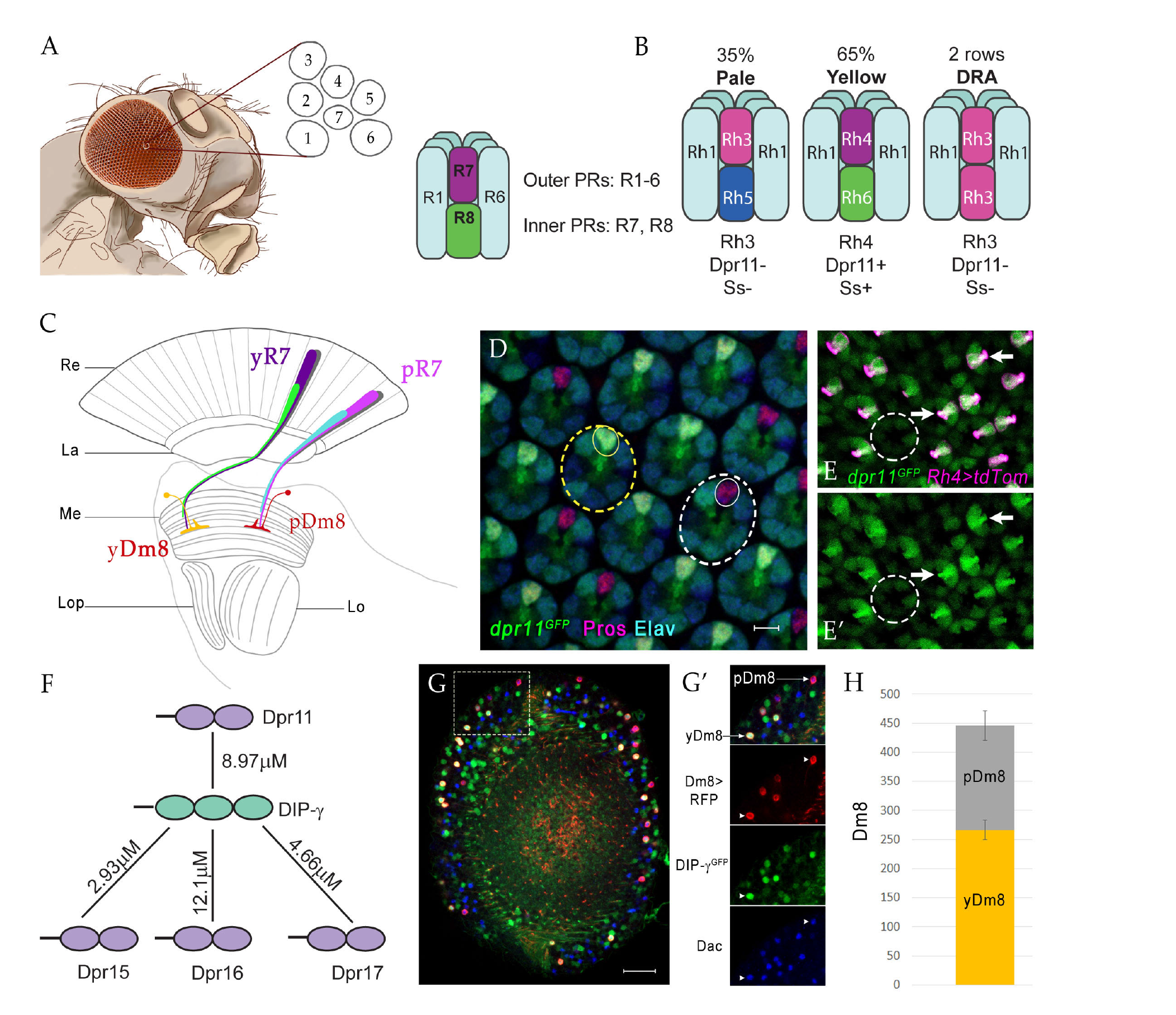
yDm8 and pDm8 populations are present in the same ratio as their presynaptic input yR7 and pR7 photoreceptors. (A-C) Overview of the Drosophila visual system. (A) Compound eye. Each ommatidium contains 8 PRs. Rhabdomeres of 7 PRs seen in diagram (R7 is stacked on top of R8). R1-6 are outer PRs, and R7 and R8 are inner PRs. (B) Three ommatidial subtypes in the retina: yellow (y), pale (p) and Dorsal Rim Area (DRA). Each ommatidium is assigned as y or p based on rhodopsin expression patterns in R7 and R8 PRs. y and p ommatidia are distributed randomly in a ∼65y:35p ratio in wild-type. Rh4, Dpr11 and Spineless transcription factor (Ss) are expressed in R7 only in y ommatidia. (C) Schematic of the adult optic lobe. yR7 and pR7, and yR8 (green) and pR8 (cyan) project to M6 and M3 layers of the medulla, respectively. The axons of outer PRs (grey) terminate in the lamina. yR7 and pR7 synapse on yDm8 and pDm8 in M6. Re: Retina; La: Lamina; Me: Medulla; Lop: Lobula plate; Lo: Lobula. (D) *dpr11^GFP^* is expressed in R7 in some ommatidia (yellow dashed circle) and absent in others (white dashed circle). R7 indicated by small circles. Mid-pupal retina labeled with anti-Pros for all R7 PRs (magenta), anti-GFP for *dpr11^GFP^* reporter (green) and anti-Elav for all neurons (blue). Maximum intensity projection; scale bar 5 µm. (E-E’) yR7 co-expresses *dpr11^GFP^* and Rh4. Dpr11 is expressed in Rh4+ R7 in yellow ommatidia (arrows) and absent from Rh3+ R7 in pale ommatidia (white dashed circle). Late pupal retina labeled with anti-RFP for Rh4>tdTomato (magenta) and anti-GFP for *dpr11^GFP^* reporter (green). (F) The DIP-γ hub consists of Dprs 11, 15, 16 and 17. Dprs are 2-IgSF domain CSPs and interact with DIPs, which are 3-IgSF domain CSPs. K_D_s shown here are from (Cosmanescu et al., 2018). Ig domains indicated by ovals. (G-G’) yDm8 and pDm8 cell bodies in adult medullary cortex. Adult optic lobes labeled with anti-RFP for pan-Dm8 driver>RFP (red), anti-GFP for *DIP-*γ*^GFP^* reporter (green) and anti-Dac for transcription factor Dachshund (blue). yDm8 expresses RFP, GFP and Dac and pDm8 expresses RFP and Dac. Inset in G shown in G’. yDm8 and pDm8 cell bodies indicated (arrows in merged, arrowheads in individual panels). Maximum intensity projection; scale bar 20 µm. (H) yDm8 and pDm8 populations are present in a 60:40 ratio, approximately matching the ratio of the input R7 PRs. The cell numbers of y and p Dm8 neurons counted with the pan-Dm8 driver and *DIP-*γ*^GFP^* are indicated on the y-axis. yDm8: 266.4+/-17.2, pDm8: 179.4+/-25.6 (n = 8-11 optic lobes; error bars indicate std. deviation).

There are two major types of ommatidia, pale (p) and yellow (y), which are randomly distributed in the retina. R7 and R8 PRs in these ommatidia are divided into subtypes with different spectral sensitivities. p ommatidia (∼35%) detect shorter wavelengths, and have R7 that express the Rh3 (shorter-wave UV) rhodopsin and R8 that express Rh5 (blue), while y ommatidia (∼65%) detect longer wavelengths, and have R7 that express Rh4 (longer-wave UV) and R8 that express Rh6 (green) (reviewed by (Viets et al., 2016))(Figure 1B). The R7 and R8 within an ommatidium mutually inhibit each other(Schnaitmann et al., 2018).

*Drosophila* phototaxes to UV (R7) in preference to visible light (R8). It also exhibits true color vision, being able to make intensity-independent discriminations among hues that differentially stimulate p and y R8 channels (Melnattur et al., 2014)(reviewed by (Song and Lee, 2018)). To be able to distinguish blue from green, or short from long-wavelength UV, the fly must have different neural responses to stimulation of p and y R8 and R7 channels and utilize these response profiles to control its actions.

In our earlier work, we showed that Dpr11 is selectively expressed by yR7 PRs (Figures 1E-E’) while its binding partner DIP-γ is expressed by a subset of Dm8 amacrine neurons, which are the primary synaptic partners for R7 (Carrillo et al., 2015). Here we show that yR7 PRs specifically connect to DIP-γ-expressing Dm8 neurons (yDm8), while pR7 PRs connect to DIP-γ-negative Dm8 neurons (pDm8) in their respective y and p “home columns”. Analysis of the electron microscopic (EM) reconstruction of the medulla (Takemura et al., 2013; Takemura et al., 2015) in light of this connection pattern shows that there are separate yellow and pale circuits that could be used for discriminating long and short-wavelength UV inputs. The yellow circuit might be constructed using DIP-γ-Dpr11 interactions, since both yDm8 and Tm5a projection neurons, which are also selectively connected to yR7 PRs, express DIP-γ (Cosmanescu et al., 2018; Karuppudurai et al., 2014). DIP-γ and Dpr11 are both required for normal morphogenesis of yDm8 distal dendrites, which fasciculate with yR7 terminals and contain many of the R7-Dm8 synapses.

Given the existence of separate yellow and pale circuits, how does the system ensure that each yR7 has a yDm8 partner? Here we show that DIP-γ-expressing yDm8 neurons are generated in excess during development and compete for presynaptic yR7 partners. yR7 PRs and yDm8 neurons recognize each other using Dpr11-DIP-γ interactions. The engagement of DIP-γ by Dpr11 is necessary for production of neurotrophic signals that allow yDm8 neurons to survive.

## RESULTS

### Dm8 and R7 subtypes are present in matching ratios

DIP-γ interacts with four Dpr partners with similar affinities: Dprs 11, 15, 16 and 17 (Figure 1F). A subset of R7 PRs in pupal retina express the *dpr11^MiMIC^* GFP reporter (henceforth denoted as *dpr11^GFP^*), and these were confirmed to be yR7 by co-labeling with a *Rh4* reporter (Figures 1B, D-E) (Carrillo et al., 2015). Reporters for Dprs 15, 16, and 17 are not detectably expressed in the pupal retina. Here we classify two subtypes of Dm8 neurons, those which express DIP-γ (yDm8) and those which do not (pDm8) (Figure 1G-G’). We determined the yDm8 population in the adult medullary cortex as those cells that express RFP under the control of a late pupal pan-Dm8 driver and GFP from the *DIP-* γ *^MiMIC^* GFP reporter (henceforth denoted as *DIP-* γ *^GFP^*). The pDm8 population was identified as those cells that express RFP but not GFP; there are no known markers or drivers that selectively label pDm8 neurons. For developmental studies, and to determine the yDm8 population without the driver, we used *DIP-*γ*^GFP^* together with the transcription factor *Dachshund (Dac),* which is expressed in both y and pDm8 neurons (Figures 1G-G’)(Hasegawa et al., 2011). Dac+, *DIP-* γ *^GFP^*+ cells are present from the beginning of pupal development.

yR7 and pR7 ommatidia are present at a 65y:35p ratio, and because of retinotopy we expect the same ratio of y and p columns in the medulla. We find that yDm8 and pDm8 neurons in wild-type are present at a ratio of 60y:40p, which is similar to the 65:35 ratio of the input R7 PRs (Figures 1B, H) (Viets et al., 2016).

Dm8 neurons are both unicolumnar and multicolumnar in their coverage of columns in the neuropil. The arbor of each Dm8 contacts 13-16 medulla columns, but most synapses are made with the R7 in the central (home) column (Fischbach and Dittrich, 1989; Gao et al., 2008; Takemura et al., 2013; Takemura et al., 2015). Figure 2A shows a horizontal view (side-view as in Figure 1C) of a rendering of an R7 terminal and aDm8 arbor from an EM reconstruction (Takemura et al., 2015). The thick bundle of dendritic processes at the center of the arbor makes extensive contacts with the R7 home column, and contains the majority of R7-Dm8 synapses (Figure 2A, Table 1). These central dendritic projections extend distally from M6 to M4, and we have denoted them as the “sprig” of a Dm8 arbor. The lateral arbor of each Dm8 overlaps extensively with other Dm8 arbors, but the center of highest arbor density, the sprig, tiles the medulla. The coverage pattern of a typical Dm8 is thus indicative of approximately 1 cell per column.

**Figure 2:**
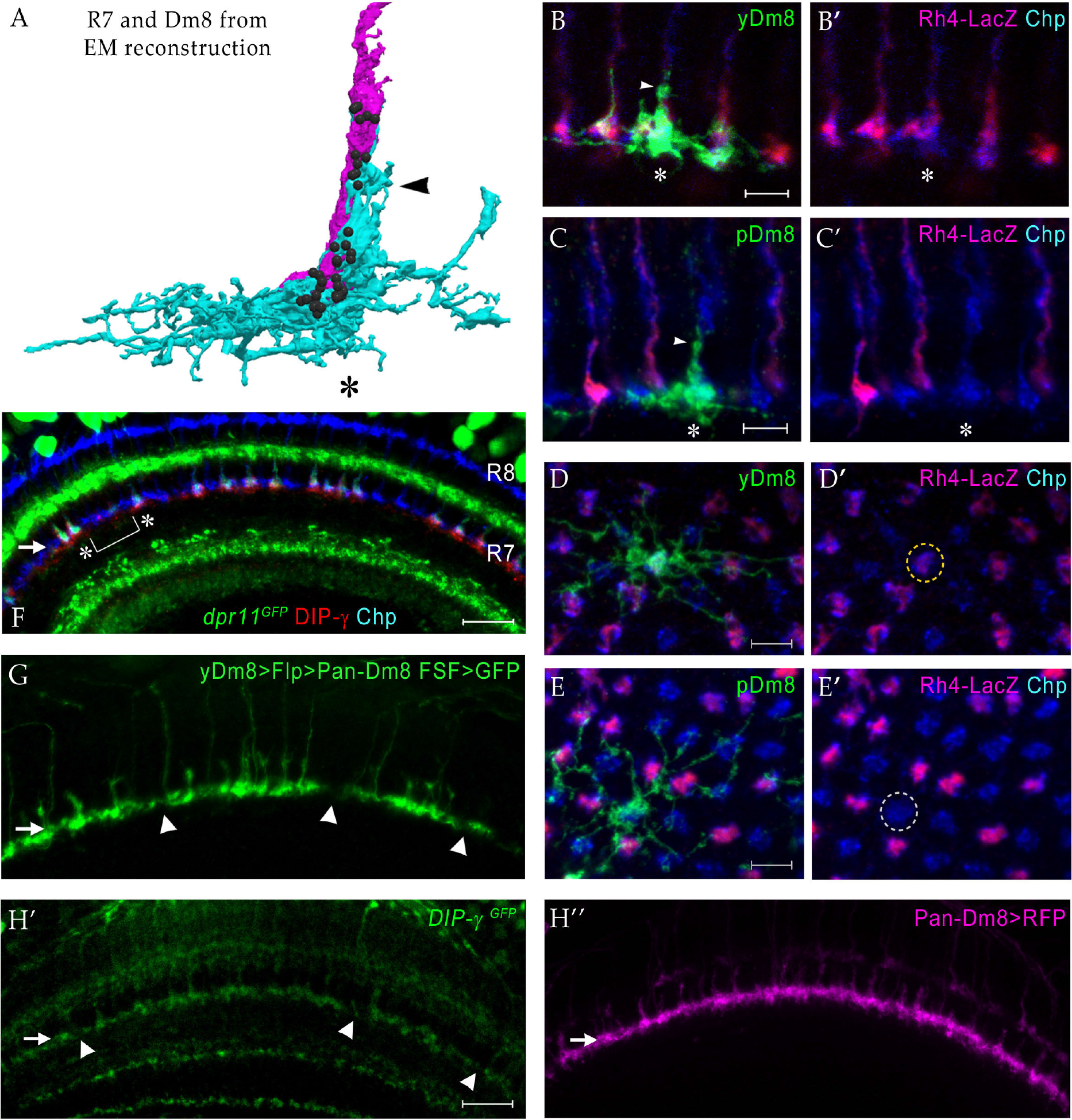
yDm8 neurons selectively innervate yR7 in their home columns and avoid pR7, and *vice versa*. (A) Rendering of a yR7 terminal and a yDm8 arbor from an EM reconstruction, in a horizontal view. The distal dendritic projection (sprig; arrowhead) of Dm8 extensively contacts the R7 home column (asterisk). Dm8, cyan; R7, magenta. Black balls indicate R7 T-bars (output synapses). (B-E) Single flipout clones generated either with the *DIP-*γ split-Gal4 Dm8 driver (denoted as yDm8 split-Gal4 driver in Figure 2–figure supplement 1A) for yDm8 or with pan-Dm8 driver for pDm8. Adult optic lobes labeled with anti-GFP for flipout clone, anti-LacZ for yR7 reporter *Rh4-lacZ* and anti-Chaoptin (Chp) for all PRs. Medulla columns were identified as pale or yellow with yR7 reporter (magenta) and Chp (blue); pR7 columns were identified by the absence of Rh4-LacZ labeling. Maximum intensity projection; scale bar 5 µm. (B-B’) Horizontal view of a yDm8. The yDm8 distal dendritic projection (sprig; arrowhead) extends distally along the home column yR7 (asterisk) to the M4 layer. (C-C’) Horizontal view of a pDm8. pDm8 has a similar morphology to yDm8, with the sprig (arrowhead) in contact with a pR7 home column (asterisk). (D-D’) Cross-sectional (top-down) view of a yDm8. The dendritic arbor of this yDm8 contacts 13 columns (9y and 4p). Home column yR7 indicated by yellow dashed circle. (E-E’) Cross-sectional view of a pDm8. The dendritic arbor of this pDm8 contacts 14 columns (10p and 5y). Home column pR7 indicated by white dashed circle. (F) DIP-γ-expressing yDm8 neurons specifically contact Dpr11-expressing yR7 home columns. Horizontal view of mid-pupal medulla labeled with anti-DIP-γ (red), anti-GFP for *dpr11^GFP^* reporter (green) and anti-Chp for all PRs (blue). All yR7 PRs have DIP-γ labeling abutting the R7 (asterisks) and none of the pR7 PRs have any DIP-γ labeling apposed to them (bracket). Maximum intensity projection; scale bar 10 µm. (G) yDm8 and pDm8 populations have independent origins. The dendritic arbors of yDm8 neurons are labeled in flies carrying DIP-γ Gal4>Flp and pan-Dm8-LexA>LexAop-FRT-stop-FRT>GFP transgenes. pDm8 that are not labeled appear as gaps (arrowheads) in the M6 layer (arrow), similar to those seen when only yDm8 neurons are labeled by *DIP-*γ*^GFP^* (H’). Adult optic lobes were labeled with anti-GFP (green). Maximum intensity projection; scale bar 10 µm. (H’-H”) Gaps representing pDm8 arbors are present in M6 layers labeled with the *DIP-*γ*^GFP^* reporter, but not in M6 labeled with the pan-Dm8 reporter. Adult optic lobes labeled with anti-GFP for *DIP-*γ*^GFP^* reporter (H’) and anti-RFP for pan-Dm8 Gal4>RFP (H”). Gaps are marked (arrowhead in H’) in the M6 layer (arrows). The pan-Dm8 driver also labels lamina neuron L3, which is seen as faint labeling above the Dm8 layer. Single confocal slice; scale bar 10 µm.

**Table 1:**
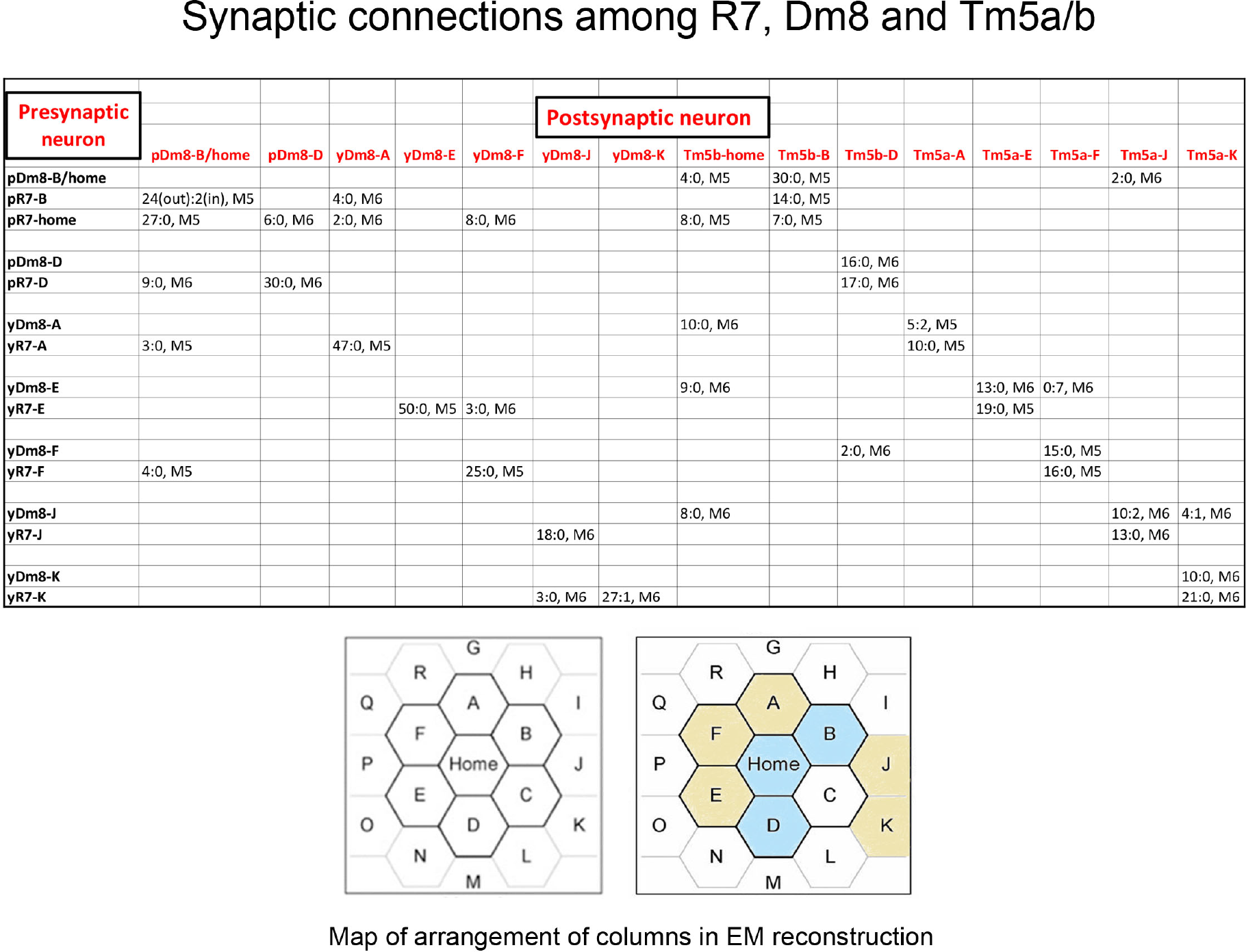
Synaptic connections among R7, Dm8, and Tm5a/b from the EM reconstruction. The entries in column1 indicate the column and y/p identities for each R7 and Dm8. The entries in row 1 indicate the column and y/p identities for the Dm8, Tm5a, and Tm5b neurons that are postsynaptic (and sometimes presynaptic) to the R7 and Dm8 neurons. In each box, the x:y nomenclature indicates the numbers of output synapses and input synapses. The layer in which most of these synapses are located is also indicated (M5 or M6). At the bottom are maps of the arrangement of columns in the reconstruction, from (Takemura et al., 2015), with y columns indicated in gold and p columns in blue in the right-hand map. We could not assign y/p identities to columns C, P, or Q, and there are no R7 outputs listed for the other columns. Tm5-C has an ambiguous morphology and could be either a Tm5a or a Tm5b, and R7-P and R7-Q have no listed synapses onto Tm5a or b. Table 1-table supplement contains additional information for non-home-column Dm8 inputs, and synapses onto Tm5c and Dm9.

Although the ratios of the input PRs and output targets match, the Dm8 cell numbers we obtained (266 yDm8 and 179 pDm8) are insufficient for innervation of the expected 730-785 yR7 plus pR7 columns in the medulla (Posnien et al., 2012) even when we account for the fact that some Dm8 neurons (∼6%; see below) have two home columns. This suggests that the Dm8 driver is not completely penetrant. Indeed, when we use Dac and *DIP-*γ*^GFP^* to determine yDm8 populations, we obtain a larger number (∼320), although this is still less than the expected total number of yDm8 neurons, which is 440-480.

### yDm8 selectively innervate yR7 in their home columns and avoid pR7

To evaluate the specificity of connections between Dm8 and R7 subtypes, we generated single-cell Dm8 (flipout) clones. For yDm8 flipouts, we used a split-GAL4 driver that includes a *DIP-*γ *^MiMIC^ GAL4-DBD* hemi-driver to selectively label DIP-γ-expressing Dm8 (Figures 2B, D, Figure 2-figure supplement 1C-C’). For pDm8 flipouts, we used a pan-Dm8 driver (Nern et al., 2015) and *Rh4-lacZ* to identify clones whose home column R7 lacked LacZ labeling, since there is no pDm8-specific driver (Figures 2C, E).

yDm8 and pDm8 arbors have similar morphologies, with the prominent dendritic sprig identifying the home column (Figures 2A-C) (Gao et al., 2008; Nern et al., 2015). Using the yDm8 split-GAL4 driver, we exclusively labeled Dm8 neurons that have yR7 as the home column (43/43 flipouts; Figures 2B, D; Figure 2-figure supplement 1A). This indicates that yDm8 specifically innervate yR7 and avoid pR7 in the home column. We also observed yDm8 with two home columns at a frequency of ∼6% (4/64 clones) (Figure 2-figure supplement 1A). All four had both sprigs on yR7 columns. Since we find that all DIP-γ-positive Dm8 neurons have yR7 home columns, we can infer that cells labeled using the pan-Dm8 driver that have pR7 home columns must be the DIP-γ-negative pDm8 subtype. Single cell clones of pDm8 were found to innervate only pR7 (21/72 total clones; one had two pR7 home columns (Figures 2C, E, Figure 2-figure supplement 1A). Specificity was further confirmed by labeling mid-pupal optic lobes with DIP-γ antibody and *dpr11*^GFP^. DIP-γ labeling is seen in the M6 layer only under yR7 that co-express Dpr11 and Chaoptin (Chp) and not under pR7 that are labeled with Chp alone (Figure 2F).

Outside of their home columns, yDm8 and pDm8 dendritic arbors contact both types of R7. Cross-sectional (top-down) views of a yDm8 clone showed that it contacted 9 yR7 and 4 pR7, while a pDm8 contacted 5 yR7 and 10 pR7 (including their respective home columns) (Figures 2D-E).

The two subtypes of Dm8 are distinguished by the expression of DIP-γ. To evaluate whether the yDm8 and pDm8 populations have separate origins, we used *DIP-*γ*-GAL4* to express FLP in a line with a flipout cassette (LexAop-FRT-stop-FRT) driving a GFP reporter and a pan-Dm8 LexA driver and examined M6 GFP labeling (Figure 2G, Figure 2-figure supplement 1B). If yDm8 and pDm8 originate as separate populations, we would expect gaps to be present in M6 (like those seen with *DIP-*γ*^GFP^* (Figure 2H’)), because the conditional *DIP-*γ GFP reporter would be expressed only in yDm8. In an alternative model, all Dm8 neurons would originate as a single DIP-γ+ population, but DIP-γ would remain on only in those Dm8 that establish a stable contact with a Dpr11+ yR7. In the subset of Dm8 that form contacts with pR7, DIP-γ would be switched off. If the latter model were correct, the conditional GFP reporter would be visualized as a continuous line in the M6 layer, similar to that seen with a pan-Dm8 driver (Figure 2H”). Figure 2G and Figure 2-figure supplement 1B show that there are gaps in the M6 layer,, indicating that the two Dm8 subtypes are in fact separate populations.

### Yellow and pale-specific synaptic connections in color vision circuits

Most R7 output synapses are made onto Dm8, Dm9 (a multicolumnar medulla intrinsic neuron), Tm5a, Tm5b, and Tm5c (Gao et al., 2008; Karuppudurai et al., 2014; Takemura et al., 2013; Takemura et al., 2015). Tm5a/b/c neurons project to the lobula, and are likely to be the main output neurons of R7 circuits. To define the synaptic connections of these neurons and determine whether they differ between the y and p columns, we examined the EM reconstruction of the medulla (Takemura et al., 2013; Takemura et al., 2015).

The morphologies of y and p R7 and Dm8 neurons, as visualized in the EM reconstruction, do not allow us to distinguish their subtypes. However, the main dendritic branch of Tm5a was found to associate with yR7 but not with pR7 axons in a light-level analysis (Karuppudurai et al., 2014), suggesting that yR7 selectively synapses onto Tm5a. Using this hypothesis as a guide, we identified five candidate yR7 PRs (in columns A, E, F, J, and K) and three pR7 PRs (in columns Home, B and D) (Table 1). There are seven Dm8 neurons whose home columns correspond to these eight R7 PRs (Dm8-B/home has two home columns). Each R7 makes many more output synapses onto its home column Dm8 than onto other Dm8 neurons, allowing us to identify B/home and D as pDm8, while A, E, F, J, and K are yDm8 (Table 1). There is no apparent specificity in synaptic connectivity between y and p R7 and Dm8 subtypes outside of the home column (Table 1-table supplement).

**Table 1:** Lines and sources:

| Genotype | Source |
| --- | --- |
| *yw hsFlp; UAS>CD2, y+>mCD8::GFP/Cyo; TM2/TM6B* | Gift from C.-H. Lee |
| *Rh4-lacZ* | Bloomington |
| *R24F06-p65.AD* | Bloomington |
| *DIP-γ Gal4-DBD* | Gift from C. Desplan |
| *OK371-VP16AD* | Gift from C.-H. Lee |
| *R24F06-Gal4* | Bloomington, (Nern et al., 2015) |
| *DIP-γ<sup>MI03222</sup>* and *DIP-γ<sup>MI03222</sup> Gal4* | H. J. Bellen |
| *DIP-γ<sup>null</sup>* | Gift from L. Zipursky, (Xu et al., 2018b) |
| *dpr11<sup>MI02231</sup>* | H. J. Bellen |
| *dpr11<sup>null</sup>* | Gift from L. Zipursky, (Xu et al., 2018b) |
| *dpr11 RNAi GD23243 (III)* | VDRC |
| *UAS-DIP-γ<sup>sh</sup> (II)* | This study |
| *UASp-DIAP I (II)* | Bloomington |
| *UAS-DIAP1-myc/TM6b* | Gift from L. Zipursky, (Xu et al., 2018b) |
| *UAS-dveA-9B2/TM3* | Gift from Hideki Nakagoshi, (Nakagawa et al., 2011) |
| *lGMR-Ss (II)* | Gift from C. Desplan |
| *lGMR-Gal4* | Bloomington |
| *sev<sup>14</sup>* | Bloomington |
| *UAS-dpr11<sup>sh</sup> (II)* | This study |
| *Rh4-lexA::p65, lexAop2-brp-short<sup>cherry</sup>* | Gift from T. Suzuki |

To analyze projection neuron input specificity, we counted synapses made by y and p R7 PRs onto Tm5a and Tm5b. The five yR7 make output synapses only onto Tm5a and not Tm5b, which was the basis for their assignment as y (Karuppudurai et al., 2014). The three pR7 synapse onto Tm5b and not Tm5a (Table 1). Thus, Tm5a and Tm5b represent separate output channels for y (Rh4) and p (Rh3) inputs for this set of columns (see Table 1-table supplement legend). Tm5a expresses DIP-γ, while Tm5b does not (Cosmanescu et al., 2018). This suggests that the selective connection of yR7 to both yDm8 and Tm5a in the home column might involve Dpr11-DIP-γ interactions. There are no GAL4 drivers that distinguish between Tm5a and Tm5b, so we cannot determine if yR7-Tm5a connections are altered in mutants.

Dm8 arbors contain both pre- and postsynaptic elements, and Dm8 is presynaptic to Tm5a/b and Dm9. We find that Dm8 output synapses onto Tm5a/b have a similar specificity to the R7 output synapses. Both pDm8 neurons almost exclusively synapse onto Tm5b and not Tm5a, and four of the five yDm8 neurons (E, F, J and K) make more synapses onto Tm5a than Tm5b (yDm8-A, however, makes ten synapses onto Tm5b-home and only five onto Tm5a-A; Table 1). Each Dm9 receives input from both types of R7 and Dm8, and makes output synapses onto multiple R7 neurons (Table 1-table supplement).

R7 synapses are polyadic, and Dm8 and Tm5a/b often sit together at synaptic sites. Figures 3F and G show R7 T-bars (active zone elements marking output synapses) adjacent to both yDm8 and Tm5a, and pDm8 and Tm5b, respectively. In column E, the yR7-E axon terminal, yDm8-E sprig, and Tm5a-E dendritic branch are tightly wrapped around each other (Figures 3A, C). yR7-E T-bars are distributed in layers M4-M6, and are apposed to postsynapses in the yDm8-E sprig and the distal dendritic branch of the Tm5a-E neuron (Figure 3C). Most yDm8-E output synapses are onto Tm5a-E and Dm9 (Table 1, Table 1-table supplement), and these are distributed between the sprig in M4-M5 and the main arbor in M6 (Figure 3C and associated videos 1 and 2). The only pDm8 with one home column in the reconstructed volume is pDm8-D. Although pDm8 usually have robust sprigs (*e.g*., Figure 2C), pDm8-D has a very thin sprig, and most pR7-D and pDm8-D T-bars are in M6 (Figure 3D and associated videos 3 and 4).

**Figure 3:**
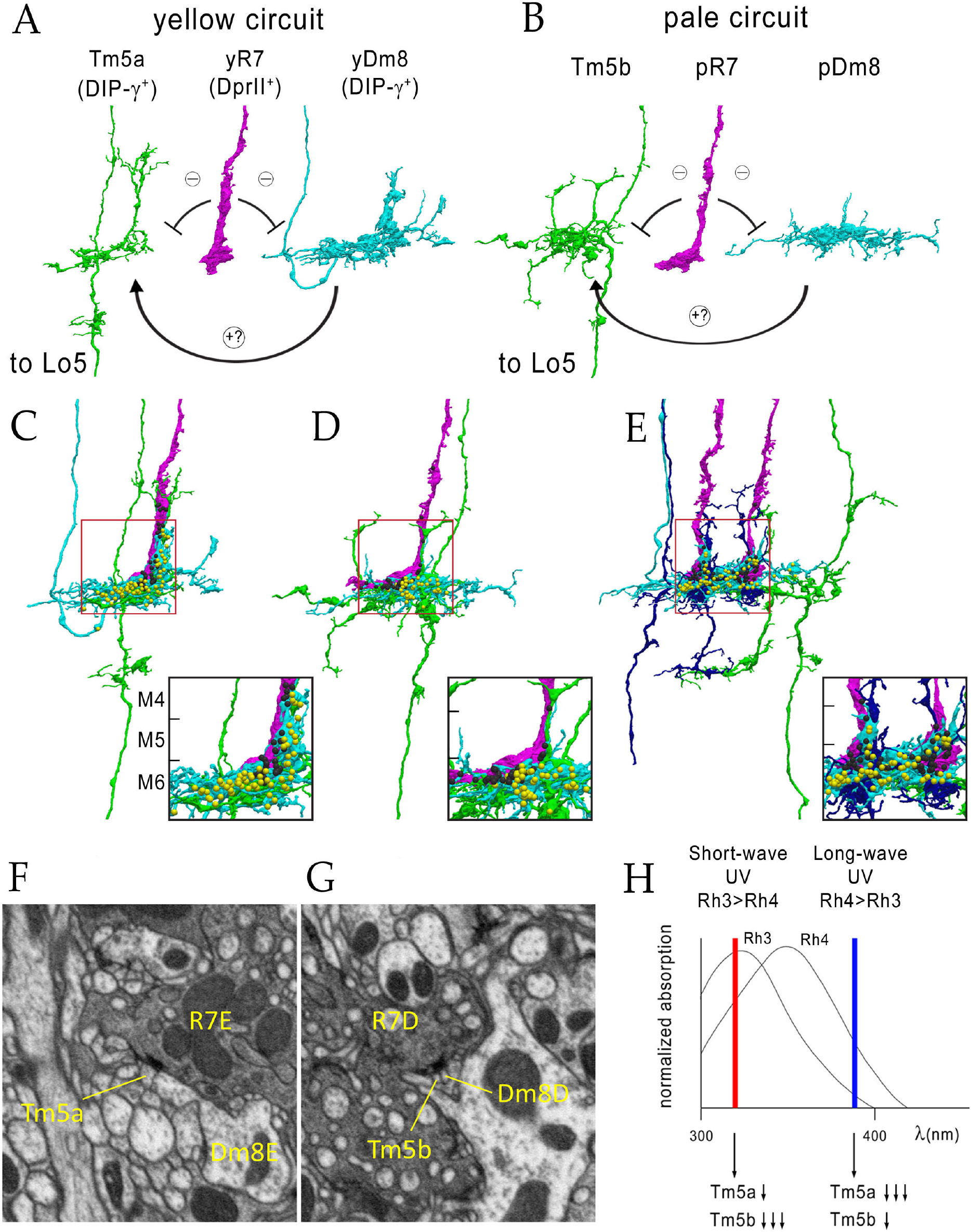
Electron microscopic reconstruction of wavelength discrimination circuits. Renderings of separated and combined cells from the EM reconstruction (Takemura et al., 2015) are shown in (A)-(E). R7, magenta; Dm8, cyan; Tm5a/b, green. (A) Cells in a yellow circuit, from column E. yR7-E, yDm8-E, and Tm5a-E are shown. yR7 inhibits both yDm8 and Tm5a (repression bars). yDm8 makes glutamatergic synapses (probably excitatory) onto Tm5a (arrow). Tm5a and Tm5b axons project to the 5^th^ layer of the lobula. (B) Cells in a pale circuit, from column D. pR7-D, pDm8-D, and Tm5b-D are shown. The connections among the cells are the same as for the yellow circuit. The implication of these connection patterns is that R7 stimulation should inhibit Tm5a/b by both direct (histaminergic) and indirect (histaminergic inhibition of excitatory Dm8 glutamatergic output) pathways. (C)-(E) Renderings of R7-Dm8-Tm5a/b circuits. The insets show the home column region where most synapses are located. Black balls, R7 T-bars; yellow balls, Dm8 T-bars. The borders of M6, M5, and M4 are indicated. (C) The column E circuit. (D) The column D circuit. (E) The column B and home circuit. This is a two-home column circuit, including pR7-B, pR7-home, pDm8-B/home, Tm5b-B, and Tm5b-home. The separated cells for this circuit are shown in Figure 3-figure supplement 1. See also associated videos 1-6 (vertical and horizontal rotation of each circuit). For a comparison of an ExM image of a wild-type yR7 and yDm8 to yR7-E and yDm8-E from the EM reconstruction, see Figure 5-figure supplement 1. (F) A section from column E, showing a polyadic synapse of yR7-E onto yDm8-E and Tm5a-E. In (F) and (G), the R7 T-bars are the black shapes on the R7 membranes where they are apposed to both postsynaptic cells. (G) A section from column D, showing a polyadic synapse of pR7-D onto pDm8-D and Tm5b-D. (H) A model for UV wavelength discrimination by the yellow and pale circuits. Short-wave UV (red bar) would stimulate Rh3+ pR7 more than Rh4+ yR7, and might therefore produce more inhibition of Tm5b than of Tm5a. Long-wave UV (blue bar) would produce more inhibition of Tm5a than of Tm5b. These signals could be read out by Lo neurons that receive Tm5a and Tm5b inputs.

pR7-B, pR7-home, pDm8-B/home, Tm5b-B, and Tm5b-home form a two-home column circuit (the individual cells of this circuit are shown in Figure 3-figure supplement 1). The B sprig of pDm8-B/home and one of the dendritic branches of Tm5b-B are both wrapped around the pR7-B terminal, while the pR7-home terminal is more loosely associated with the home sprig of pDm8-B/home and dendritic branches of both Tm5b neurons (Figure 3E and associated videos 5 and 6, and Figure 3-figure supplement 1). All pR7-B→Tm5b synapses (14) and most pDm8-B/home→Tm5b synapses (30 *vs.* 4) are onto Tm5b-B. pR7-home makes similar numbers of synapses onto Tm5b-B and Tm5b-home (Table 1).

### *DIP-*γ and *dpr11* mutations cause abnormalities in yDm8 dendritic arbors

If the yellow circuit is constructed using DIP-γ-Dpr11 interactions, as suggested by our analysis of the EM reconstruction, one might expect that there would be abnormalities in yDm8 arbors in *DIP-*γ and *dpr11* mutants. To assess yDm8 arbor morphology in whole-animal mutants, we surveyed horizontal views of the neuropil under conditions where we genetically converted all Dm8 neurons to yDm8 (see below) (Figures 4B-D). Both mutants displayed atypical patterns when labeled with the pan-Dm8 driver, with changes in the regular pattern of sprigs extending upward along the home column R7.

**Figure 4:**
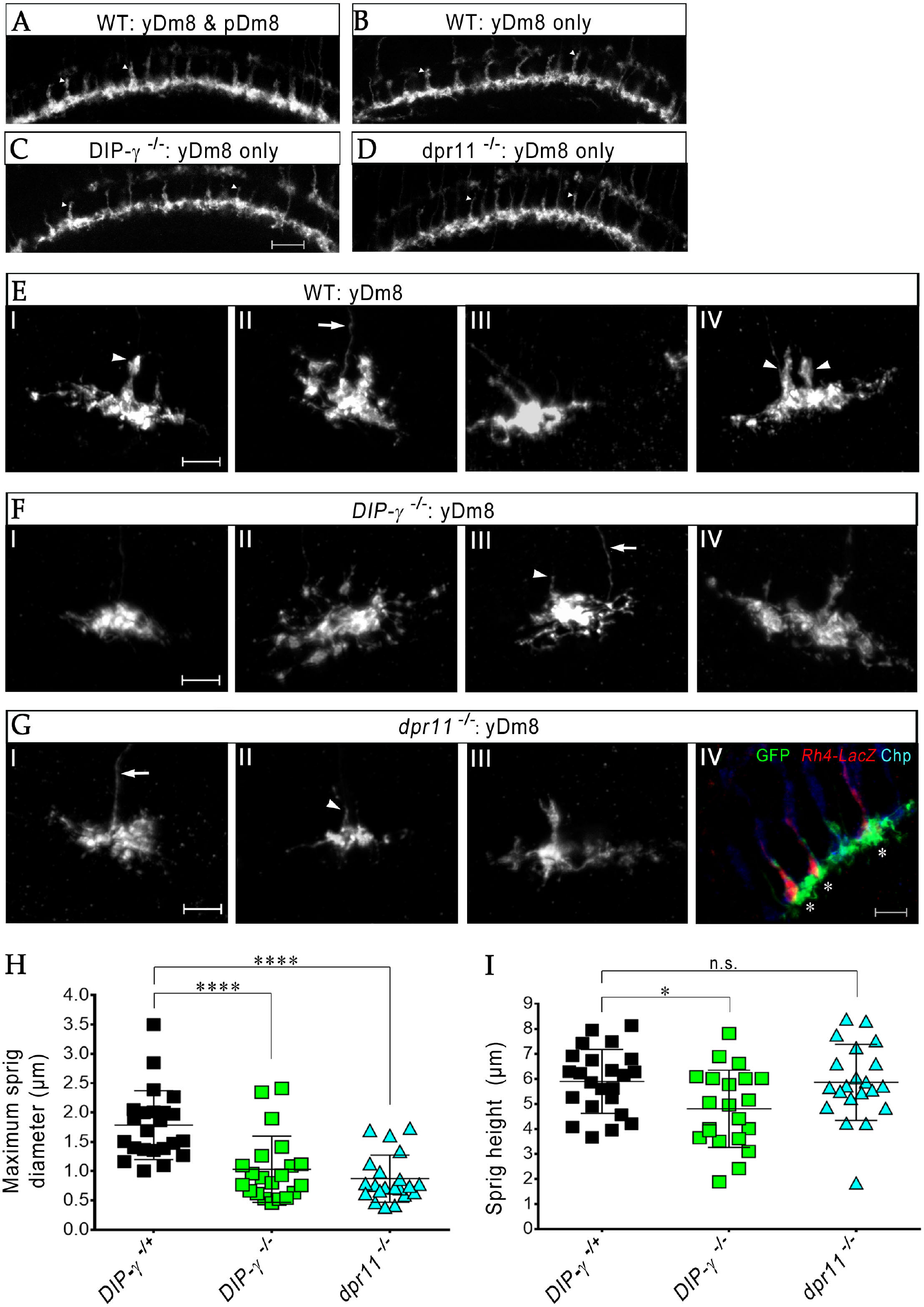
Loss of DIP-γ or Dpr11 alters morphology of yDm8 distal dendrites. (A-D) Horizontal views of the M6 layer labeled with the pan-Dm8 driver in (A) WT, (B) *ss^GOF^*, (C) *ss^GOF^*; *DIP-*γ *^-/-^*, (D) *ss^GOF^*; *dpr11^-/-^*. In *ss^GOF^* (B-D), all Dm8 are converted to yDm8 (see Figure 8). The pan-Dm8 driver also labels lamina neuron L3, which is seen as faint labeling above the Dm8 layer. Sprigs (arrowheads) are thinner or are missing in (C) and (D) as compared to (B). Maximum intensity projection; scale bar 10 µm. (E-G) yDm8 flipouts generated in (E) wild-type, (F) *DIP-*γ mutant and (G) *dpr11* mutant. Variations observed in yDm8 morphology for each genotype shown. Sprigs marked with arrowheads in panels E-I, E-IV, F-III, and G-II. Axons marked with arrows in E-II, F-III, and G-I. Panel E-IV is a 2-home column yDm8 with both sprigs marked. Flipouts in wild-type and *DIP-*γ mutants were generated with the yDm8 split-Gal4 driver; *dpr11* mutant flipouts were generated with the pan-Dm8 driver and scored as yDm8 using Rh4-LacZ labeling (red in panel G-IV; yDm8 clones labeled with asterisks). Quantitation of clones for all three genotypes in Figure 4-figure supplement 1. Maximum intensity projection; scale bar 5 µm. (H-I) Maximum sprig diameter is reduced significantly in both *dpr11* and *DIP-*γ mutants. Sprig height was slightly affected in *DIP-*γ mutants but not in *dpr11* mutants. Graph shows mean +/-std. deviation and unpaired Student’s t-test p-values. Sprig diameter: *DIP-*γ *^-/^*^+^ 1.8+/-0.6, *DIP-*γ *^-/^*^-^ 1.0+/-0.6, *dpr11 ^-/-^* 0.9+/-0.4, ****p<0.0001 for both mutants. Sprig height: *DIP-*γ *^-/^*^+^ 5.9+/-1.3, *DIP-*γ *^-/^*^-^ 4.8+/-1.5, *dpr11 ^-/-^* 5.9+/-1.5, from left to right *p=0.014, not significant (n.s.)

We generated flipouts in wild-type and mutants (Figure 4-figure supplement 1A) and examined arbor morphology (Figures 4E-G). The sprig, which is located in the center of the arbor, usually has an expanded region at the distal end in M4/M5 (see Figures 2A-C). There is a large variation in sprig diameter and height in both wild-type and mutants. Maximum sprig diameter was significantly decreased in both *DIP-*γ and *dpr11* mutants compared to control, but the height of the sprig was relatively unaffected (Figures 4H-I).

### yDm8 arbor morphology in detail

To obtain high-resolution views of interactions between yDm8 sprigs and R7 terminals, we examined single cell yDm8 clones in wild-type and *DIP-*γ mutant brains using expansion microscopy (ExM)(Figure 5 and associated videos 1-4) ((Mosca et al., 2017); reviewed by (Karagiannis and Boyden, 2018)). The morphology of the arbor in M6 and of the sprig in M4 and M5 can be visualized in detail in ExM images (Figure 5A). The sprig is on a home column R7, where most of the synapses from yR7 to yDm8 are located (Figures 5B-C, Figure 5-figure supplement 1). In addition to the sprig, this yDm8 arbor has two thin dendritic processes that emerge from the base and are wrapped around neighboring non-home column R7 terminals (Figure 5C). The home column R7 extends into the base of the sprig, making multiple contacts with the base of the Dm8 arbor. *DIP-*γ mutant yDm8 neurons have thinner sprigs, in agreement with our quantitative analysis of mutant flipout clones (Figures 4H, 5D; Figure 5-figure supplement 2B).

**Figure 5:**
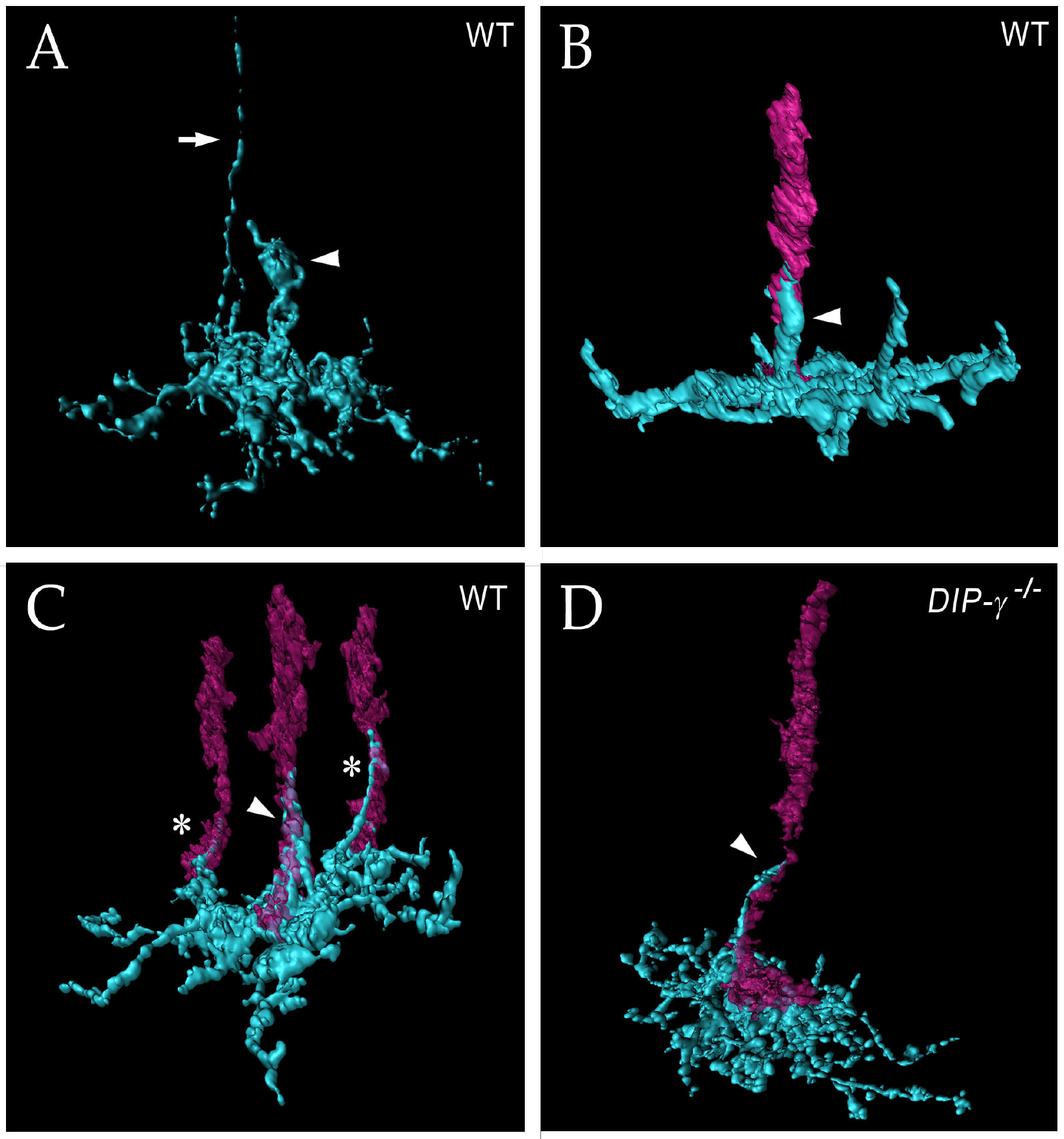
yDm8 arbor morphology in wild-type and *DIP-*γ mutant visualized with expansion microscopy. yDm8 dendritic arbors (cyan) visualized with expansion microscopy and surface rendered with Imaris software. (A-C) Wild-type; (D) DIP-γ *^-/-^*. Arrowheads, sprigs; arrow in (A), axon. In (B)-(D), one or more R7 terminals/axons are included in the rendering. Two different views of the same flipout clone are shown in panels (B) and (C). The R7 terminals/axons are semi-transparent in (C) and (D). The home column R7 located at the center of the arbor makes extensive contacts with the sprig as well as with the base of the dendritic arbor in M6 in (B) and (C). Two thinner dendritic processes positioned on the edges of the arbor (asterisks) contact two non-home column R7 in (C). The yDm8 in the *DIP-*γ *^-/-^* mutant has a much thinner sprig as compared to wild-type (D). See associated videos 1-4 (vertical and horizontal rotations) and Figure 5-supplemental figure 2 for additional views of the wild-type clone in (C) and the mutant clone in (D).

**Table 2:** List of genotypes in figures and graphs:

| Short genotype | Complete genotype | Figures |
| --- | --- | --- |
| *dpr11<sup>GFP</sup>/+* | *dpr11<sup>MI02231-GFP</sup>/+* | 1D |
|  | *Rh4 Gal4; dpr11<sup>MI02231-GFP</sup>/10XUAS-tdTomato::myr* | 1E-E' |
|  | *10xUAS-mCD8::RFP/+; DIP-γ<sup>MI03222</sup>/R24F06-Gal4* | 1G-G', H |
| yDm8 split-Gal4 | *R24F06-p65.AD; DIP-γ<sup>MI03222</sup>Gal4.DBD* | 2B, D, 4E-F |
| Pan-Dm8 driver | *R24F06-Gal4 (III)* | 2C, E, 4G |
|  | *yw hsflp/+; UAS>CD2, y+>mCD8::GFP/R24F06-p65.AD; Rh4-lacZ/DIP-γ<sup>MI03222</sup>Gal4.DBD* | 2B, D |
|  | *yw hsflp/+; UAS>CD2, y+>mCD8::GFP/+; Rh4LacZ/R24F06-Gal4* | 2C, E |
|  | *dpr11<sup>MI02231-GFP</sup>/+* | 2F |
| yDm8>Flp>Pan-Dm8 FSF>GFP | *20xUAS-flp/+; R24F06-LexA/+; DIP-γ<sup>MI03222</sup>Gal4, LexAop FRT>stop>FRT mCD8::GFP/+* | 2G |
|  | *10xUAS mCD8::RFP/+; DIP-γ<sup>MI03222-GFP</sup>/R24F06-Gal4* | 2H'-H'' |
| WT | *10xUAS-mCD8::RFP/+; DIP-γ<sup>MI03222-GFP</sup>/R24F06Gal4* | 4A |
| WT | *IGMR-ss/10xUAS-mCD8::RFP; DIP-γ<sup>MI03222-GFP</sup>/R24F06Gal4* | 4B |
| *DIP-γ<sup>-/-</sup>* | *IGMR-ss/10xUAS-mCD8::RFP; DIP-γ<sup>MI03222-GFP</sup>/R24F06Gal4, DIP-γ null* | 4C |
| *ss<sup>GOF</sup>; dpr11<sup>-/-</sup>* | *IGMR-ss/10xUAS-mCD8::RFP; R24F06-Gal4, dpr11<sup>null</sup>/dpr11<sup>null</sup>, DIP-γ<sup>MI03222-GFP</sup>* | 4D |
| *DIP-γ<sup>-/+</sup>* | *yw hsflp/+; UAS>CD2, y+>mCD8::GFP/R24F06-p65.AD; Rh4-lacZ/DIP-γ<sup>MI03222</sup>Gal4.DBD* | 4E, H-I |
| *DIP-γ<sup>-/-</sup>* | *yw hsflp/+; UAS>CD2, y+>mCD8::GFP/R24F06-p65.AD; DIP-γ null/DIP-γ<sup>MI03222</sup>Gal4.DBD* | 4F, H-I |
| *dpr11<sup>-/-</sup>* | *yw hsflp/+; UAS>CD2, y+>mCD8::GFP/Rh4LacZ; R24F06-Gal4, dpr11<sup>null</sup>/dpr11<sup>null</sup>* | 4G-I |
| WT | *yw hsflp/+;* | 5A-C & Videos |
|  | *UAS>CD2, y+>mCD8::GFP/R24F06-p65.AD;*<br>*Rh4-lacZ/DIP-$\gamma^{MI03222}$Gal4.DBD* | 1-2 |
| *DIP-$\gamma^{-/-}$* | *yw hsflp/+; UAS>CD2, y+>mCD8::GFP/R24F06-p65.AD;*<br>*DIP-$\gamma^{null}$/DIP-$\gamma^{MI03222}$Gal4.DBD* | 5D & Videos 3-4 |
| *DIP-$\gamma^{-/+}$* | *10xUAS-mCD8::RFP/+; R24F06-Gal4/ DIP-$\gamma^{MI03222}$-GFP* | 6A-B, D |
| *DIP-$\gamma^{-/-}$* | *10xUAS-mCD8::RFP/+;*<br>*R24F06-Gal4, DIP-$\gamma^{null}$/DIP-$\gamma^{MI03222}$-GFP* | 6A-B |
| *dpr11<sup>-/+</sup> control* | *10xUAS-mCD8::RFP/+;*<br>*R24F06-Gal4, dpr11<sup>null</sup>/DIP-$\gamma^{MI03222}$-GFP* | 6A-B |
| *dpr11<sup>-/-</sup>* | *10xUAS-mCD8::RFP/+;*<br>*R24F06-Gal4, dpr11<sup>null</sup>/ dpr11<sup>null</sup>, DIP-$\gamma^{MI03222}$-GFP* | 6A-B, H |
| *DIP-$\gamma^{-/+}$ control* | *R24F06-LexA/+;*<br>*13xLexAop-tdTomato::myr, DIP-$\gamma^{MI03222}$-GFP /+* | 6A-B |
| *DIP-$\gamma^{-/-}$ control* | *R24F06-LexA/+; DIP-$\gamma^{MI03222}$ Gal4/*<br>*13xLexAop-tdTomato::myr, DIP-$\gamma^{MI03222}$-GFP* | 6A-B, E |
| *DIP-γ rescue* | *UAS-DIP-$\gamma^{sh}$/R24F06-LexA; DIP-$\gamma^{MI03222}$ Gal4/*<br>*13xLexAop-tdTomato::myr, DIP-$\gamma^{MI03222}$-GFP* | 6A-B, F |
| *yDm8 DIAP rescue* | *UASp-DIAP/R24F06-LexA; DIP-$\gamma^{MI03222}$ Gal4/*<br>*13xLexAop-tdTomato::myr, DIP-$\gamma^{MI03222}$-GFP* | 6A-B, G |
| *DIP-$\gamma^{-/+}$* | *DIP-$\gamma^{MI03222}$-GFP /+* | 6C |
| *DIP-$\gamma^{-/-}$* | *DIP-$\gamma^{MI03222}$-GFP / DIP-$\gamma^{null}$* | 6C |
| *dpr11<sup>-/-</sup>* | *dpr11<sup>null</sup>/dpr11<sup>null</sup>, DIP-$\gamma^{MI03222}$-GFP* | 6C |
| *lGMR Gal4 control* | *lGMR-Gal4/+; dpr11<sup>null</sup>, DIP-$\gamma^{MI03222}$-GFP /+ @29deg* | 6C |
| *lGMR Gal4/dpr11 RNAi* | *lGMR-Gal4/+; dpr11<sup>null</sup>, DIP-$\gamma^{MI03222}$-GFP / dpr11 RNAi*<br>*GD23243 (III) @29deg* | 6C |
| *lGMR Gal4/+* | *lGMR-Gal4/+; DIP-$\gamma^{MI03222}$-GFP /+* | 6C |
| *lGMR Gal4/ UAS-DIP-γ* | *lGMR-Gal4/UAS-DIP-γ; DIP-$\gamma^{MI03222}$-GFP /+* | 6C |
| *DIP-γ*<sup>-/+</sup> | *DIP-γ*<sup>MI03222-GFP/+</sup> | 7A, B-D |
| *DIP-γ*<sup>-/-</sup> | *DIP-γ*<sup>MI03222-GFP</sup> / *DIP-γ*<sup>null</sup> | 7A |
|  | *dpr11*<sup>MI02231-GFP</sup> | 7E |
|  | WT control | 7F |
| WT | *dpr11*<sup>MI02231-GFP/+</sup> | 8A |
| *ss*<sup>GOF</sup> | *lgmr-ss/+*; *dpr11*<sup>MI02231-GFP/+</sup> | 8B |
| *DIP-γ*<sup>-/+</sup> | *10xUAS-mCD8::RFP/+*; *R24F06-Gal4* / *DIP-γ*<sup>MI03222-GFP</sup> | 8C-D |
| *LGMR>ss*; *DIP-γ*<sup>-/+</sup> | *10xUAS-mCD8::RFP* / *lgmr-ss*;<br>*DIP-γ*<sup>MI03222-GFP</sup> / *R24F06-Gal4* | 8C-D |
| *LGMR>ss*; *DIP-γ*<sup>-/-</sup> | *10xUAS-mCD8::RFP* / *lgmr-ss*;<br>*DIP-γ*<sup>MI03222-GFP</sup> / *R24F06-Gal4*, *DIP-γ*<sup>null</sup> | 8C-D |
| *DIP-γ*<sup>-/-</sup> | *10xUAS-mCD8::RFP/+*;<br>*DIP-γ*<sup>MI03222-GFP</sup> / *R24F06-Gal4*, *DIP-γ*<sup>null</sup> | 8C-D |
| *sev*<sup>-/+;;</sup> *DIP-γ*<sup>-/+</sup> | *sev*<sup>14/+</sup>; ; *R24F06-Gal4* / *UAS-mCD8::RFP*, *DIP-γ*<sup>MI03222-GFP</sup> | 8C-D |
| *sev*<sup>-/-;;</sup> *DIP-γ*<sup>-/+</sup> | *sev*<sup>14/sev</sup><sup>14</sup>; ;<br>*R24F06-Gal4* / *UAS-mCD8::RFP*, *DIP-γ*<sup>MI03222-GFP</sup> | 8C-D |
| *sev*<sup>-/-;;</sup> *DIP-γ*<sup>-/-</sup> | *sev*<sup>14/sev</sup><sup>14</sup>; ;<br>*R24F06-Gal4*, *DIP-γ*<sup>null</sup> / *UAS-mCD8::RFP*, *DIP-γ*<sup>MI03222-GFP</sup> | 8C-D |
| *lgmr Gal4* control | *R24F06-LexA/+*;<br>*13xLexAop-tdTomato::myr*, *DIP-γ*<sup>MI03222-GFP</sup> / *lgmr-Gal4* | 8C-D |
| *lgmr Gal4>*<br>*UAS-dpr11* | *UAS-Dpr11-sh* / *R24F06-LexA*;<br>*13xLexAop-tdTomato::myr*, *DIP-γ*<sup>MI03222-GFP</sup> / *lgmr-Gal4* | 8C-D |
| WT | *lgmr- Gal4/+* | 8E |
| *dpr11*<sup>GOF</sup> | *lgmr-Gal4* / *UAS-dpr11*<sup>sh</sup> | 8F |
| *lgmr Gal4* control | *lgmr-Gal4/+*; *DIP-γ*<sup>MI03222-GFP/+</sup> | 8G |
| *lGMR Gal4>*<br>*UAS-Dve* | *lGMR-Gal4/+; DIP-$\gamma^{M103222-GFP}$ / UAS-dveA-9B2* | 8G |
| *lGMR Gal4* | *lGMR- Gal4/+* | 8H |
| *lGMR Gal4>*<br>*UAS-dpr11* | *lGMR-Gal4/UAS-dpr11<sup>sh</sup>* | 8H |

**Table 3:** List of genotypes in Supplemental figures and graphs:

| Short genotype | Complete genotype | Supplemental figures |
| --- | --- | --- |
| yDm8 split-Gal4 driver | *yw hsflp/+;*<br>*UAS>CD2, y+>mCD8::GFP/ R24F06-p65.AD;*<br>*Rh4-lacZ/DIP-$\gamma^{M103222}Gal4.DBD$* | Figure 2-figure supplement 1A |
| Pan-Dm8 driver | *yw hsflp/+; UAS>CD2, y+>mCD8::GFP/+;*<br>*Rh4LacZ/R24F06-Gal4* | Figure 2-figure supplement 1A |
| yDm8>Flp>Pan-Dm8 FSF>GFP | *20xUAS-flp/+; R24F06-LexA/+;*<br>*DIP-$\gamma^{M103222}Gal4$, LexAop FRT>stop>FRT*<br>*mCD8::GFP/+* | Figure 2-figure supplement 1B |
|  | *yw hsflp/+;*<br>*UAS>CD2, y+>mCD8::GFP/OK371 dVP16.AD;*<br>*Rh4 LacZ/DIP-$\gamma^{M103222}Gal4.DBD$* | Figure 2-figure supplement 1C-C' |
| WT:<br>Pan-Dm8 driver | *yw hsflp/+; UAS>CD2, y+>mCD8::GFP/+;*<br>*Rh4LacZ/ R24F06-Gal4* | Figure 4-figure supplement 1A |
| WT:<br>yDm8 split-Gal4 driver | *yw hsflp/+;*<br>*UAS>CD2, y+>mCD8::GFP/R24F06-p65.AD;*<br>*Rh4-lacZ/DIP-$\gamma^{M103222}Gal4.DBD$* | Figure 4-figure supplement 1A |
| *DIP-$\gamma^{-/-}$ :*<br>yDm8 split -Gal4 driver | *yw hsflp/+;*<br>*UAS>CD2, y+>mCD8::GFP/R24F06-p65.AD;*<br>*DIP-$\gamma^{null}$ /DIP-$\gamma^{M103222}Gal4.DBD$* | Figure 4-figure supplement 1A |
| *dpr11<sup>-/-</sup> :* | *yw hsflp/+; UAS>CD2, y+>mCD8::GFP/Rh4LacZ;* | Figure 4-figure |
| Pan-Dm8 driver | *R24F06-Gal4, dpr11<sup>null</sup>/dpr11<sup>null</sup>* | supplement 1A |
| WT | *yw hsflp/+;*<br>*UAS>CD2, y+>mCD8::GFP/R24F06-p65.AD;*<br>*Rh4-lacZ/DIP-γ<sup>M103222</sup>Gal4.DBD* | Figure 5-figure<br>supplement 1A,<br>Figure 5-figure<br>supplement 2A |
| *DIP-γ<sup>-/-</sup>* | *yw hsflp/+;*<br>*UAS>CD2, y+>mCD8::GFP/R24F06-p65.AD;*<br>*DIP-γ<sup>null</sup>/DIP-γ<sup>M103222</sup>Gal4.DBD* | Figure 5-figure<br>supplement 2B |
| *DIP-γ<sup>GFP</sup>, DIP-γ*<br>*Gal4>UAS-DIAP<sup>myc</sup>* | *UAS-diap1.myc, DIP-γ<sup>M103222-GFP</sup>/*<br>*DIP-γ<sup>M103222</sup>Gal4* | Figure 6-figure<br>supplement 1A |
|  | *DIP-γ<sup>M103222-GFP</sup>/+* | Figure 7-figure<br>supplement 1A-A' |
|  | WT control | Figure 7-figure<br>supplement 1B-B' |
|  | *dpr11<sup>M102231-GFP</sup>/+* | Figure 7-figure<br>supplement 1C |
| *ss<sup>GOF</sup>* | *lGMR-ss/+; DIP-γ<sup>M103222-GFP</sup>* | Figure 8-figure<br>supplement 1A |
| *ss<sup>GOF</sup>; DIP-γ<sup>-/-</sup>* | *lGMR-ss/+; DIP-γ<sup>M103222-GFP</sup>/DIP-γ<sup>null</sup>* | Figure 8-figure<br>supplement 1B |
| WT | *lGMR-Gal4/+; DIP-γ<sup>M103222-GFP</sup>/+* | Figure 8-figure<br>supplement 1C |
| *dpr11<sup>GOF</sup>* | *lGMR-Gal4/UAS-dpr11<sup>sh</sup>; DIP-γ<sup>M103222-GFP</sup>/+* | Figure 8-figure<br>supplement 1D |
| *lGMR-Gal4>UAS-Dve* | *lGMR-Gal4/+; DIP-γ<sup>M103222-GFP</sup>/UAS-dveA-9B2* | Figure 8-figure<br>supplement 1E |
| *sev<sup>-/-</sup>* | *sev<sup>14</sup>/sev<sup>14</sup>; R24F06Gal4/DIPgMiMIC, UAS-mCD8RFP* | Figure 8-figure<br>supplement 1F |
| WT | *lGMR Gal4/+* | Figure 8-figure<br>supplement 1G |
| *lGMR-Gal4>UAS-dpr11* | *lGMR Gal4/+; UAS-dpr11<sup>sh</sup>* | Figure 8-figure supplement 1H |
| *dpr11<sup>GOF</sup>* | *lGMR Gal4/+; UAS-dpr11<sup>sh</sup>* | Figure 8-figure supplement 1I |
| *ss<sup>GOF</sup>* | *lGMR-ss/+* | Figure 8-figure supplement 1J |
| WT | *dpr11<sup>MI02231-GFP/+</sup>* | Figure 8-figure supplement 2A-A' |
| *dpr11<sup>-/-</sup>* | *dpr11<sup>MI02231-GFP/dpr11<sup>null</sup></sup>* | Figure 8-figure supplement 2B-B' |
| *dpr11<sup>+/-</sup>* | *dpr11<sup>MI02231-GFP/+</sup>* | Figure 8-figure supplement 2C |
| *dpr11<sup>-/-</sup>* | *dpr11<sup>MI02231-GFP/dpr11<sup>null</sup></sup>* | Figure 8-figure supplement 2C |
| *dpr11<sup>+/-</sup>* | *Rh4-lexA::p65, lexAop2-brp-short<sup>cherry</sup>/+;*<br>*dpr11<sup>null</sup>/+* | Figure 8-figure supplement 2D |
| *dpr11<sup>-/-</sup>* | *Rh4-lexA::p65, lexAop2-brp-short<sup>cherry</sup>/+;*<br>*dpr11<sup>null</sup>/dpr11<sup>null</sup>* | Figure 8-figure supplement 2D |

### Interaction between Dpr11 in yR7 and DIP-γ in yDm8 is required for yDm8 survival

We previously showed that Dm8 cells are lost in *DIP-*γ *^GFP^/Df* animals (Carrillo et al 2015). To explore this in detail, we used CRISPR-generated null mutants for both *DIP-*γ and *dpr11* (Xu et al., 2018b) and determined yDm8 and pDm8 populations (Figures 6A-C). *DIP-*γ *^GFP^*, which has no detectable protein expression, was used as one of the *DIP-*γ alleles to allow identification of yDm8 soma. Both *DIP-*γ and *dpr11* mutants showed a ∼50% decrease in yDm8 cell number using the pan-Dm8 driver and 60-65% loss when determined with Dac (Figures 6A, C). The loss of yDm8 in *DIP-*γ mutants was partially rescued by expressing a *DIP-*γ transgene using the *DIP-*γ *Gal4* driver (Figure 6A). To determine if yDm8 loss in *dpr11* mutants was due to the absence of Dpr11 from the eye or from other Dpr11*-*expressing cells in the medulla, we performed eye-specific *dpr11* transgenic RNAi, and found a ∼40% reduction in yDm8 cell numbers (Figure 6C).

**Figure 6:**
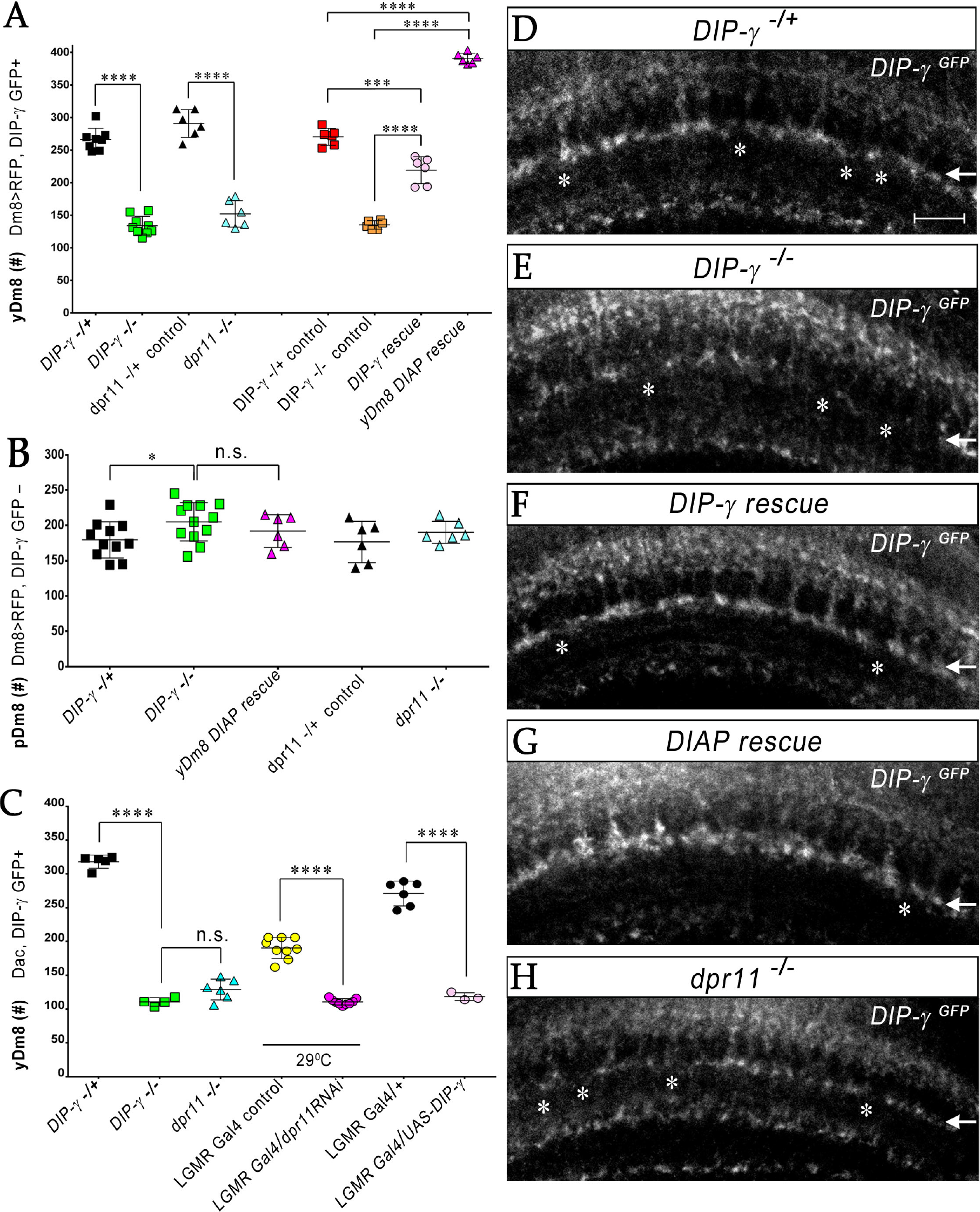
yR7 PRs provide neurotrophic support to yDm8 neurons through Dpr11-DIP-γ interactions. (A-B) yDm8 and pDm8 cell numbers were determined using pan-Dm8 driver>RFP and *DIP-* γ*^GFP^* reporter. yDm8 neurons express both RFP and GFP and pDm8 neurons express only RFP. (C) yDm8 cell number determined using anti-GFP for *DIP-*γ*^GFP^* reporter and anti-Dac. (A-C) *DIP-*γ*^GFP^* reporter (indicated in all graphs as *DIP-*γ ***^-^****^/+^*) is an insertion in the 5’ UTR intron that has no detectable protein expression. Thus, this line serves as a mutant as well as a reporter of *DIP-*γ transcript expression. Graphs show mean +/-std. deviation and unpaired Student’s t-test p-values. Complete genotypes in Materials and Methods (Table 2). (A) Both *DIP-*γ and *dpr11* mutants show ∼50% loss of yDm8 neurons, and this is rescued in *DIP-*γ mutants by inhibiting cell death with DIAP. yDm8 cell numbers in heterozygous controls, *DIP-*γ and *dpr11* mutants, and *DIP-*γ and DIAP rescues in *DIP-*γ mutant are shown. *DIP-*γ *^-/^*^+^ 266.4+/-17.2, *DIP-*γ *^-/^*^-^ 134+/-14.3, ****p<0.0001; *dpr11 ^-/+^* control 290.7+/-21.3, *dpr11 ^-/-^* 152+/-20.4, ****p<0.0001; *DIP-*γ *^-/^*^+^ control 270+/-13.03, *DIP-*γ rescue 219.2+/-20.7, ***p=0.0004; *DIP-*γ *^-/^*^-^ mutant control 135.2+/-6.5, *DIP-*γ rescue 219.2+/-20.7, ****p<0.0001; *DIP-*γ *^-/^*^+^ control 270+/-13.03, *DIAP* rescue 390.8+/-8.0, ****p<0.0001; *DIP-*γ *^-/^*^-^ mutant control 135.2+/-6.5, *DIAP* rescue 390.8+/-8.0, ****p<0.0001. (B) pDm8 cell numbers in both mutants and in DIAP rescue. *DIP-*γ *^-/^*^+^ 179.4+/-25.6, *DIP-*γ *^-/^*^-^ 204.9+/-27.2, *p=0.03; *dpr11 ^-/+^* control 176.5+/-29.2, *dpr11 ^-/-^* 190.2+/-15.5, not significant (n.s.) p=0.34; *DIP-*γ *^-/^*^-^ 204.9+/-27.2, *DIAP* yDm8 rescue 191.8+/-23.2, not significant (n.s.) p=0.32 (C) Dpr11 in R7 is required for yDm8 survival. yDm8 cell number in wild-type, mutants, dpr11 eye-specific RNAi, and ectopic DIP-γ expression in PRs. *DIP-*γ *^-/+^* 318+/-9.7, *DIP-*γ *^-/^*^-^ 110.5+/-6.1, ****p<0.0001; *DIP-*γ *^-/^*^-^ 110.5+/-6.1, *dpr11 ^-/-^* 129+/-15.4, not significant (n.s.) p=0.055; *lGMR-Gal4* control at 29^0^C 190.3+/-15.5, *lGMR-Gal4>dpr11 RNAi* 110.6+/-4.5 ****p<0.0001 *lGMR-Gal4* control 217+/-18.3, *lGMR-Gal4>UAS-DIP-*γ 118.3+/-5.9 ****p<0.0001 (D-H) yDm8 labeling in the neuropil in wild-type, *DIP-*γ mutant, *DIP-*γ rescue, *DIAP* yDm8 rescue, and *dpr11* mutant, using *DIP-*γ*^GFP^* reporter. Large gaps (asterisks) representing yDm8 cell death are seen in the M6 layer (arrow) in both *DIP-*γ and *dpr11* null mutants (E, H), whereas wild-type, *DIP-*γ and *DIAP* rescues showed smaller and fewer gaps (asterisks in D, F-G). Adult optic lobes were labeled with anti-GFP for yDm8 reporter.

We also ectopically expressed DIP-γ in PRs, and found that this phenocopied the *DIP-*γ and *dpr11* loss-of-function (LOF) phenotypes (Figure 6C). This result suggests that the presence of both Dpr11 and its partner on the same cells (expression in *cis*) prevents Dpr11 in yR7 from interacting with DIP-γ in *trans* (on yDm8). Similar results were observed for DIP-α and Dpr10 in the neuromuscular system, where DIP-α is normally expressed on the RP2 motoneuron and Dpr10 on its muscle targets. LOF mutants lacking either protein are missing specific muscle branches, and ectopic expression of DIP-α on muscles, or of Dpr10 on RP2, generates the same phenotypes (Ashley et al., 2019).

To assess whether the decrease in yDm8 population was due to apoptotic cell death, we expressed *Drosophila* inhibitor of apoptosis protein 1 (DIAP) in *DIP-*γ expressing cells (Figure 6-figure supplement 1). DIAP overexpression rescued the loss of yDm8 in *DIP-*γ mutants, indicating that DIP-γ is required for cell survival and that yDm8 undergo apoptosis when it is absent (Figure 6A). Furthermore, the number of yDm8 that survived when DIAP was expressed was ∼45% greater than the number of yDm8 present in controls. It has been reported that extensive cell death occurs in the medullary cortex in wild-type during normal optic lobe development (Togane et al., 2012). Thus, in addition to the yDm8 that were rescued from cell death caused by the absence of DIP-γ, DIAP also rescued yDm8 that were lost due to normal developmental cell death (see Discussion). The fact that *dpr11* and *DIP-*γ mutants displayed a similar extent of yDm8 loss implies that Dpr11 in yR7 signals to yDm8 *via* DIP-γ to ensure their survival, implicating yR7 as a source of neurotrophic support for yDm8. Absence of either molecule compromises the interaction and leads to death of yDm8 neurons.

We next assessed pDm8 populations in the above genotypes and found that in general they did not differ significantly from controls. Importantly, pDm8 cell numbers did not decrease when the yDm8 population was increased by DIAP rescue (Figure 6B; see below).

We also visualized the M6 layer using the *DIP-*γ *^GFP^* reporter, and found that morphologies of the yDm8 layer in the above genotypes were consistent with the results obtained in the cortex. In the neuropil, larger gaps were observed in the mutants, and fewer and smaller gaps in wild-type and rescue genotypes (Figures 6D-H).

*DIP-*γ mutants also displayed a ∼4.5-fold higher frequency of two-home column flipouts as compared to control (Figure 4-figure supplement 1). The increase in two-home column yDm8 may be a response strategy in which the surviving yDm8 innervate more yR7 in order to compensate for the loss of neurotrophic support mediated by DIP-γ.

### DIP-γ controls yDm8 death by interacting with Dpr11 during early pupal development

To determine when the Dpr11 and DIP-γ interaction is needed for yDm8 survival, we examined the timecourse of yDm8 loss in wild-type and *DIP-*γ mutants starting at 15h after puparium formation (APF) (Figure 7A). In wild type, the population of yDm8 is unchanged between 15h APF and 25h APF. yDm8 cell numbers in *DIP-*γ mutants are similar to wild-type cell numbers at 15h APF, but there is significant cell death by 25h APF. By 45h APF, the yDm8 population in both wild type and mutant had reached the adult cell number. *DIAP* rescue of the *DIP-*γ mutant restored the population in the adult back to the original pool size at 15h APF (Figure 7A). Thus, DIP-γ is required for suppression of yDm8 cell death early in pupal development. Extensive migration of yDm8 cells was observed in wild-type, and this migration was unaffected in the *DIP-*γ mutant (Figures 7B-D, and data not shown).

**Figure 7:**
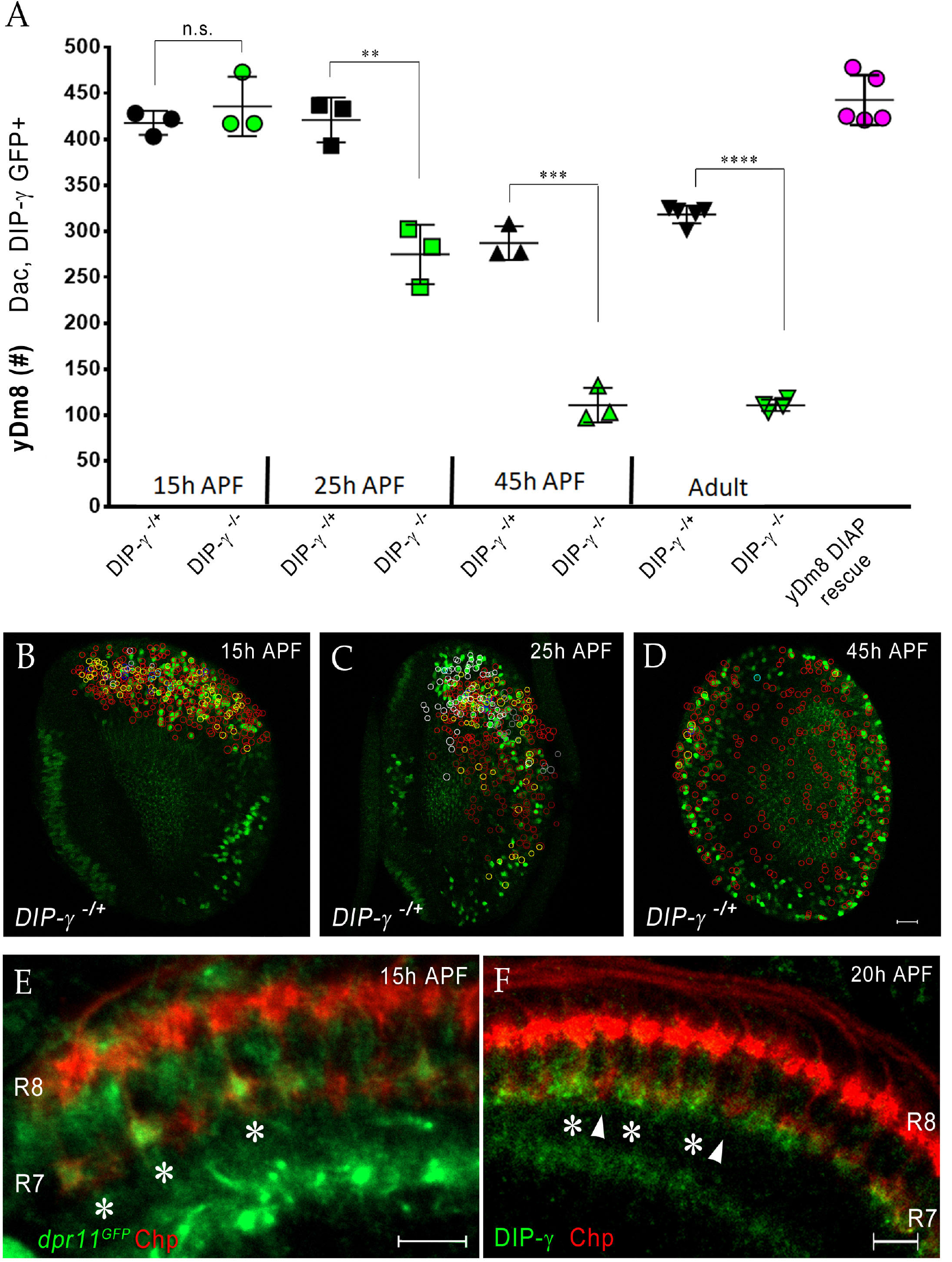
Dpr11-DIP-γ interactions are required early in pupal development to prevent yDm8 cell death. (A) yDm8 cell death occurs in *DIP-*γ mutants between 15h and 45h APF. yDm8 cell death in wild-type occurs between 25h and 45h APF. DIAP expression in the *DIP-*γ mutant rescues cell number back to the original level at 15 APF. yDm8 cell number determined with anti-GFP for *DIP-*γ *^GFP^* reporter and anti-Dac. *DIP-*γ*^GFP^* reporter heterozygote indicated as *DIP-*γ ***^-^****^/+^*. Graph shows mean +/-std. deviation and unpaired Student’s t-test p-values. Complete genotypes in Materials and Methods (Table 2). 15h APF: *DIP-*γ *^-/+^* 417.7+/-13.1, *DIP-*γ *^-/^*^-^ 435.7+/-32.3, not significant (n.s.) p= 0.4218; 25h APF: *DIP-*γ *^-/+^* 421+/-24.3, *DIP-*γ *^-/^*^-^ 274.7+/-32.3, **p= 0.0033; 45h APF: *DIP-*γ *^-/+^* 287+/-18.2, *DIP-*γ *^-/^*^-^ 110.7+/-18.7, ***p=0.0003; 25h APF *DIP-*γ *^-/+^* 421+/-24.3, 45h APF *DIP-*γ *^-/+^* 287+/-18.2, **p=0.0016; Adult: *DIP-*γ *^-/+^* 318+/-9.7, *DIP-*γ *^-/^*^-^ 110.5+/-6.1, ****p<0.0001; yDm8 *DIAP* rescue 442.6+/-27.2, 15h APF *DIP-*γ *^-/^*^+^ 417.7+/-13.1, not significant (n.s.) p=0.20. (B-D) yDm8 cell bodies migrate during early pupal development. yDm8 cell bodies labeled with anti-Dac (not shown here) and anti-GFP for *DIP-*γ *^GFP^* reporter are circled to indicate position in the cortex in wild-type at 15h, 25h and 45h APF. There is no difference in their relative positions in *DIP-*γ mutants as compared to that in wild-type (data not shown). (E) Dpr11 is expressed in select R7 PRs at 15h APF (asterisks). *dpr11^GFP^* reporter labeled with anti-Chp (red) and anti-GFP (green) at 15h APF. Note that one of the younger R7 PRs located on the right of the image also expresses Dpr11. Single confocal slice; scale bar 5 µm. (F) DIP-γ is expressed in yDm8 neurons apposed to specific R7 PRs by 20h APF (asterisks). pR7 terminals (arrowheads) do not show overlapping DIP-γ labeling (see also Figure 7-figure supplement 1B-B’). Wild-type, labeled at 20h APF with anti-DIP-γ (green) and anti-Chp (red). Single confocal slice; scale bar 10 µm.

In order for the interaction to occur, yR7 and its target yDm8 have to be in close proximity, and Dpr11 and DIP-γ need to be expressed. We found that *dpr11* was expressed in select R7 at 15h APF, suggesting that the yellow fate of R7 was determined by that time (Figure 7E). We next examined *DIP-*γ expression in early pupa and found DIP-γ labeling adjacent to select R7 at 20h APF, indicating that yDm8 arbors are apposed to yR7 terminals (Figure 7F). The gaps in DIP-γ labeling are presumed to be pDm8 (Figure 7-figure supplement 1). Taken together, our results are consistent with a model in which selection of yDm8 by yR7 occurs by 15h-20h APF *via* Dpr11-DIP-γ binding and thereby ensures survival of yDm8.

### Perturbing R7 fate in the retina alters Dm8 fate in the medulla

Since Dpr11 is expressed in yR7 PRs, we next examined how the two subtypes of Dm8 neurons respond when R7 fates are altered. The transcription factor Spineless (Ss) controls yellow and pale subtype choice in the retina. Ectopic expression of *ss* in all PRs (*ss^GOF^*) induces yellow fate in all R7 cells and prevents formation of pR7 (Wernet et al., 2006). We first determined Dpr11 and DIP-γ expression in *ss^GOF^*. In contrast to wild-type, where only yR7 is labeled by *dpr11^GFP^,* all R7 terminals showed GFP expression when *ss* was mis-expressed (Figures 8A-B). Thus, Ss expression activates Dpr11 along with Rh4 and other yR7-specific genes. The normal gaps in *DIP-*γ*^GFP^* labeling in the M6 layer were missing in *ss^GOF^*. When *DIP-*γ was removed in the *ss^GOF^* background, gaps were again observed in the yDm8 layer, indicating that the M6 signal in *ss^GOF^* is indeed due to DIP-γ-expressing yDm8 (Figure 8-figure supplement 1A-C). The change in R7 fate is thus transmitted to the downstream circuit in the medulla *via* Dpr11-DIP-γ interactions, resulting in uniform yDm8 labeling in the M6 layer.

**Figure 8:**
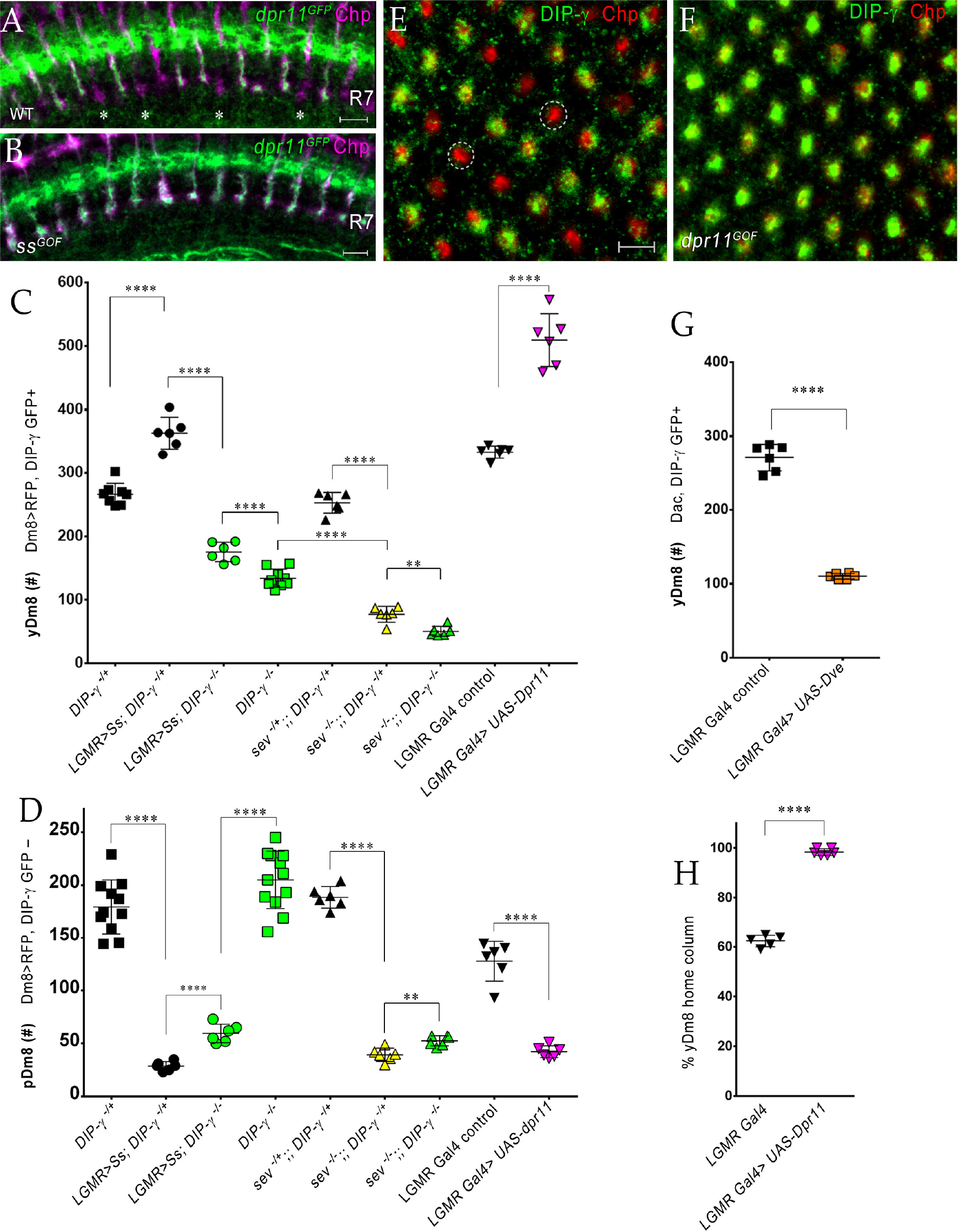
The representation of Dm8 subtypes in the medulla is altered by manipulation of R7 fates in the retina. (A-B) Conversion of all R7 PRs to the yR7 fate in *ss^GOF^* results in Dpr11 expression in every R7. Dpr11 is expressed in yR7 PRs in (A) wild-type and (B) *ss^GOF^*. pR7 PRs are present in wild-type (asterisk) and absent in *ss^GOF^*. Mid-pupal optic lobes labeled with anti-GFP (green) for *dpr11^GFP^* reporter and anti-Chp (magenta) for all PRs. Maximum intensity projection; scale bar 5 µm. (C-D) yDm8 and pDm8 cell number determined using pan-Dm8 driver>RFP and *DIP-*γ*^GFP^* reporter. yDm8 expresses both RFP and GFP and pDm8 expresses only RFP. Graphs show mean +/-std. deviation and unpaired Student’s t-test p-values. Complete genotypes in Materials and Methods (Table 2). *DIP-*γ*^GFP^* reporter heterozygote indicated as *DIP-*γ ***^-^****^/+^*. (C) More yDm8 neurons survive when all R7 PRs are converted to the yR7 fate (*ss^GOF^*) or when they all express Dpr11 (*dpr11^GOF^*). yDm8 neurons are also lost when R7 PRs are absent (*sev*) and/or *DIP-*γ is mutant. yDm8 cell number in wild-type, *ss^GOF^*, *sev* mutant, *dpr11^GOF^*: *DIP-*γ *^-/^*^+^ 266.4+/-17.2, *ss^GOF^*, *DIP-*γ *^-/^*^+^ 363+/-25.3, ****p<0.0001; *ss^GOF^*, *DIP-*γ *^-/^*^+^ 363+/-25.3, *ss^GOF^*, *DIP-*γ *^-/^*^-^ 175.3+/-15.2, ****p<0.0001; *DIP-*γ *^-/^*^-^ 134+/-14.3, *ss^GOF^*, *DIP-*γ *^-/^*^-^ 175.3+/-15.2,, ***p=0.0001 *sev ^-/+^*;; *DIP-*γ *^-/^*^+^ 252.8+/-16.3, *sev ^-/-^*;; *DIP-*γ *^-/^*^+^ 77.3+/-12.5, ****p<0.0001 *sev ^-/-^*;; *DIP-*γ *^-/+^* 77.3+/-12.5, *sev ^-/-^*;; *DIP-*γ *^-/-^* 50.5+/-7.8, **p=0.0012 *DIP-*γ *^-/-^* 134+/-14.3*, sev ^-/-^*;; *DIP-*γ *^-/-^* 50.5+/-7.8, ****p<0.0001 *lGMR Gal4* control 333+/-9.8*, lGMR Gal4>UAS-dpr11* 509.7+/-41.5, ****p<0.0001 (D) pDm8 cell numbers decrease dramatically when R7 PRs are absent, when they are converted to the yR7 fate (*ss^GOF^*), or when they all express Dpr11 (*dpr11^GOF^*). pDm8 cell number in wild-type, *ss^GOF^*, *sev* mutant, Dpr11 overexpression: *DIP-*γ *^-/^*^+^ 179.4+/-25.6, *ss^GOF^*, *DIP-*γ *^-/^*^+^ 28.7+/-4.3, ****p<0.0001; *ss^GOF^*, *DIP-*γ *^-/^*^+^ 28.7+/-4.3, *ss^GOF^*, *DIP-*γ *^-/^*^-^ 59.5+/-8.8, ****p<0.0001; *DIP-*γ *^-/^*^-^ 204.9+/-27.2, *ss^GOF^*, *DIP-*γ *^-/^*^-^59.5+/-8.8, ****p<0.0001; *lGMR Gal4* control 127.7+/-18.7, *lGMR Gal4>UAS-dpr11* 42.2+/-5.6, ****p<0.0001 *sev ^-/+^*;; *DIP-*γ *^-/^*^+^ 187.4+/-11.0, *sev ^-/-^*;; *DIP-*γ *^-/^*^+^ 39.2+/-7.0, ****p<0.0001 *sev ^-/-^*;; *DIP-*γ *^-/+^* 39.2+/-7.0, *sev ^-/-^*;; *DIP-*γ *^-/-^* 51.8+/-5.0, **p=0.005 (E-F, H) Dpr11 overexpression in the retina converts all medulla columns to y by selecting for yDm8. Pupal medullary neuropil (∼45h-48hAPF) of (E) *lGMR-Gal4* control and (F) *lGMR-Gal4>UAS-dpr11* labeled with anti-Chp (red) and anti-DIP-γ (green). Cross-section views of the medulla shown (E-F). Quantitation in (H). Maximum intensity projection; scale bar 5 µm. (G) Conversion of all R7 PRs to pR7 fate results in loss of yDm8 neurons. yDm8 cell number in *dve^GOF^* counted with anti-Dac and anti-GFP for *DIP-*γ*^GFP^* reporter. *lGMR Gal4* control 271+/-18.3, *lGMR Gal4>UAS-Dve* 110.3+/-3.8, ****p<0.0001. Graph shows mean +/-std. deviation and unpaired Student’s t-test p-values. (H) Percentage of Dm8 home columns that are yDm8 (quantitated from images like those in (E) and (F)). *lGMR Gal4/+* 0.62+/-0.02 (n=199), *lGMR Gal4>UAS-dpr11* 0.98+/-0.01 (n=234), ****p<0.0001; n represents total number of columns analyzed. Graph shows mean +/-std. deviation and unpaired Student’s t-test p-values.

We determined yDm8 and pDm8 cell number in the above genotypes (Figures 8C-D). In *ss^GOF^*, we observed a 36% increase in yDm8, while pDm8 cell number was decreased by 84%. The increase in yDm8 in *ss^GOF^* was dependent on DIP-γ, as absence of *DIP-*γ in the *ss^GOF^* background resulted in 53% loss of yDm8 neurons. Interestingly, pDm8 increased significantly, doubling their cell numbers as compared to *ss^GOF^* alone (Figure 8D). Thus, yDm8 and pDm8 populations compete with each other when R7 fate is altered, as observed in *ss^GOF^*.

To assess Dm8 subtypes in the reverse situation where there are only pR7 in the eye, we ectopically expressed the Defective proventriculus (Dve) transcription factor, a repressor downstream of *ss*, in all PRs (Johnston et al., 2011; Nakagawa et al., 2011; Yan et al., 2017). We used this strategy because *ss* LOF mutations cause lethality and we observed no phenotype with eye-specific *ss* RNAi. We predicted that there would be fewer yDm8 neurons in *dve^GOF^*, as there are no yR7 to provide neurotrophic support. Since the external eye morphologies of these animals were abnormal, we initially examined the neuropil layers of the medulla. Except for the *DIP-*γ*^GFP^* labeling in M6, other layers of the neuropil showed similar patterns to wild-type. The M6 layer had large gaps, similar to those observed in *DIP-*γ mutants, suggesting that there were fewer yDm8 neurons (Figure 8-figure supplement 1E). Indeed, yDm8 cell number decreased by ∼60% as compared to the control (Figure 8G).

Sevenless is a receptor tyrosine kinase that is required for specification of R7 in the developing eye. In *sev* mutants, R7 is absent, because the R7 precursor cell becomes a non-neuronal cone cell. We found that both the yDm8 and pDm8 populations were reduced in *sev* mutants by >70% as compared to control cell numbers, showing that both classes of Dm8 depend on R7 PRs for neurotrophic support. yDm8 cell numbers were further reduced when DIP-γ was removed in the *sev* mutant background (Figure 8C). This dependence on DIP-γ likely occurs because there are a few Dpr11-expressing R7 remaining in the *sev* mutant (Figure 8-figure supplement 1F), although the allele we used is described as an amorph. The ∼50 yDm8 that remain in a *sev DIP-*γ double mutant may not require trophic support for survival, or might obtain support through a different pathway. pDm8 numbers are slightly increased by removal of DIP-γ in the *sev* mutant background. This is probably because they are competing with yDm8 for the few remaining R7 PRs, so that the loss of additional yDm8 through removal of DIP-γ frees up those slots to be occupied by pDm8.

In order to complete our analysis of the consequences of changing R7 fates, we examined ectopic expression of Dpr11 in the eye (*dpr11^GOF^*). We reasoned that since *dpr11* was expressed in all R7 in *ss^GOF^* (Figure 8B), Dpr11 overexpression in the eye should mimic *ss^GOF^*. Indeed, the M6 layer showed continuous yDm8 labeling with no pDm8 gaps, similar to *ss^GOF^* (Figure 8-figure supplement 1D). This was confirmed by determining yDm8 and pDm8 cell numbers in *dpr11^GOF^*. The yDm8 population increased by 53% over control when Dpr11 was overexpressed in all PRs, accompanied by a 67% loss of pDm8 (Figures 8C-D). Dpr11 overexpression in *dpr11^GOF^* was confirmed by labeling with Dpr11 antibody. Dpr11 overexpression does not convert pR7 to the yR7 fate, because R7 PRs that ectopically express Dpr11 do not all express Rh4 (Figure 8-figure supplement 1G-I).

We next examined cross-sectional (top-down) views of the mid-pupal medulla in *dpr11^GOF^* labeled with DIP-γ and Chp antibodies to assess the composition of Dm8 columns when Dpr11 is expressed in all R7 PRs. Wild-type controls showed 62y:38p columns in the medulla, similar to the 65y:35p ratio of yellow and pale ommatidia in the eye. By contrast, in *dpr11^GOF^* only yellow columns were observed, indicating that all pDm8 home columns had been replaced by yDm8 (Figures 8E-F). Thus∼35% of yDm8 neurons had now selected PRs that express Dpr11 but were otherwise of the p subtype as their home column R7 (Figure 8H, Figure 8-figure supplement 1I). Similar results were obtained in *ss^GOF^*, in which pR7 PRs are actually converted to the y subtype (Figure 8-figure supplement 1J).

### yDm8 do not mistarget to pR7 in *dpr11* mutants

Since ectopic expression of Dpr11 in all PRs changed the identities of home column Dm8, converting them all to yDm8 (Figures 8E-F, H), we investigated whether there were defects in home column selection in *dpr11* mutants. We examined mutants and heterozygous controls at 44h APF because Dm8 have not yet contacted neighboring columns at this stage, making it possible to unambiguously assign the home column (Ting et al., 2014). y and p R7 were identified using *dpr11^GFP^* and Chp antibody. Using the DIP-γ antibody to label yDm8, we determined how many yR7 containing columns were paired with yDm8. In controls, every yR7 column had DIP-γ labeling adjacent to it. In *dpr11* mutants, there was a significantly higher percentage of unpaired yR7 as compared to controls (Figure 8-figure supplement 2C). However, this result can be explained by the fact that yDm8 die in *dpr11* mutants, so that the number of surviving yDm8 neurons is insufficient to innervate all yR7. Thus, the absence of DIP-γ labeling adjacent to yR7 does not indicate that Dpr11 is required for home column selection. We found no mistargeting errors where pR7 containing columns labeled only with Chp were apposed to yDm8 in *dpr11* mutants (Figure 8-figure supplement 2A-B). However, pR7 may not be available as targets because they would have been selected by pDm8 neurons. The fact that pR7 are occupied by pDm8 in mutants makes it difficult to assess whether Dpr11-DIP-γ interactions are required for selection of yDm8 as targets by yR7.

## DISCUSSION

A circuit for UV wavelength discrimination in *Drosophila* is defined by expression of a pair of cell-surface IgSF binding partners, Dpr11 and DIP-γ. Dpr11 is selectively expressed by Rh4+ yR7 PRs (Figure 1), which connect to the yDm8 subtype of Dm8 amacrine neurons that express DIP-γ. Rh3+ pR7 connect to the pDm8 subtype, which is DIP-γ negative (Figure 2). yR7 and yDm8 synapse onto Tm5a neurons, which also express DIP-γ and project to the lobula (Figure 3 and associated videos), while pR7 and pDm8 synapse onto Tm5b. Dpr11-DIP-γ interactions are also involved in determination of Dm8 arbor morphology (Figures 4, 5 and associated videos). yDm8 neurons are generated in excess during development and compete for yR7 partners. Their survival is controlled by neurotrophic signaling mediated by transsynaptic Dpr11-DIP-γ interactions (Figures 6, 7). yDm8 and pDm8 neurons do not normally compete for survival signals, but can be forced to do so by changing R7 subtype fate (Figure 8).

### Control of R7-Dm8 connectivity by neurotrophic Dpr11-DIP-γ signaling

Yellow and pale ommatidia are generated by a stochastic process and are distributed in a ∼65y:35p ratio in the retina (Wernet et al., 2006). Since yDm8 and pDm8 are separate populations (Figure 2) and R7 y *vs.* p fates are randomly determined, what mechanisms ensure that each yDm8 has a yR7 partner and each pDm8 has a pR7 partner? The strategy appears similar to those used in the development of many mammalian nervous system circuits, in that yDm8 neurons are generated in excess of their final cell numbers in adults, and those that are not selected for survival by an appropriate home column R7 partner die through apoptosis.

Our conclusions are summarized in a model (Figure 9). There are two subtypes of Dm8: yDm8 that express DIP-γ and pDm8 that do not. yDm8 are born in excess of the final cell numbers that are present in the adult (Figure 7A). There are likely to be extra pDm8 as well, but due to the lack of a pDm8 marker we were unable to determine their cell numbers in early pupae. Dpr11, the binding partner of DIP-γ, is expressed exclusively in yR7 in the retina (Figures 1D-E), allowing yDm8 neurons to select yR7 PRs as their appropriate partners. The extra yDm8 that do not find a yR7 partner undergo cell death due to lack of neurotrophic support. This selection mechanism ensures that the ratio of yDm8 to pDm8 in the medulla matches the ratio of yR7 to pR7 in the eye (Figure 1H).

**Figure 9:**
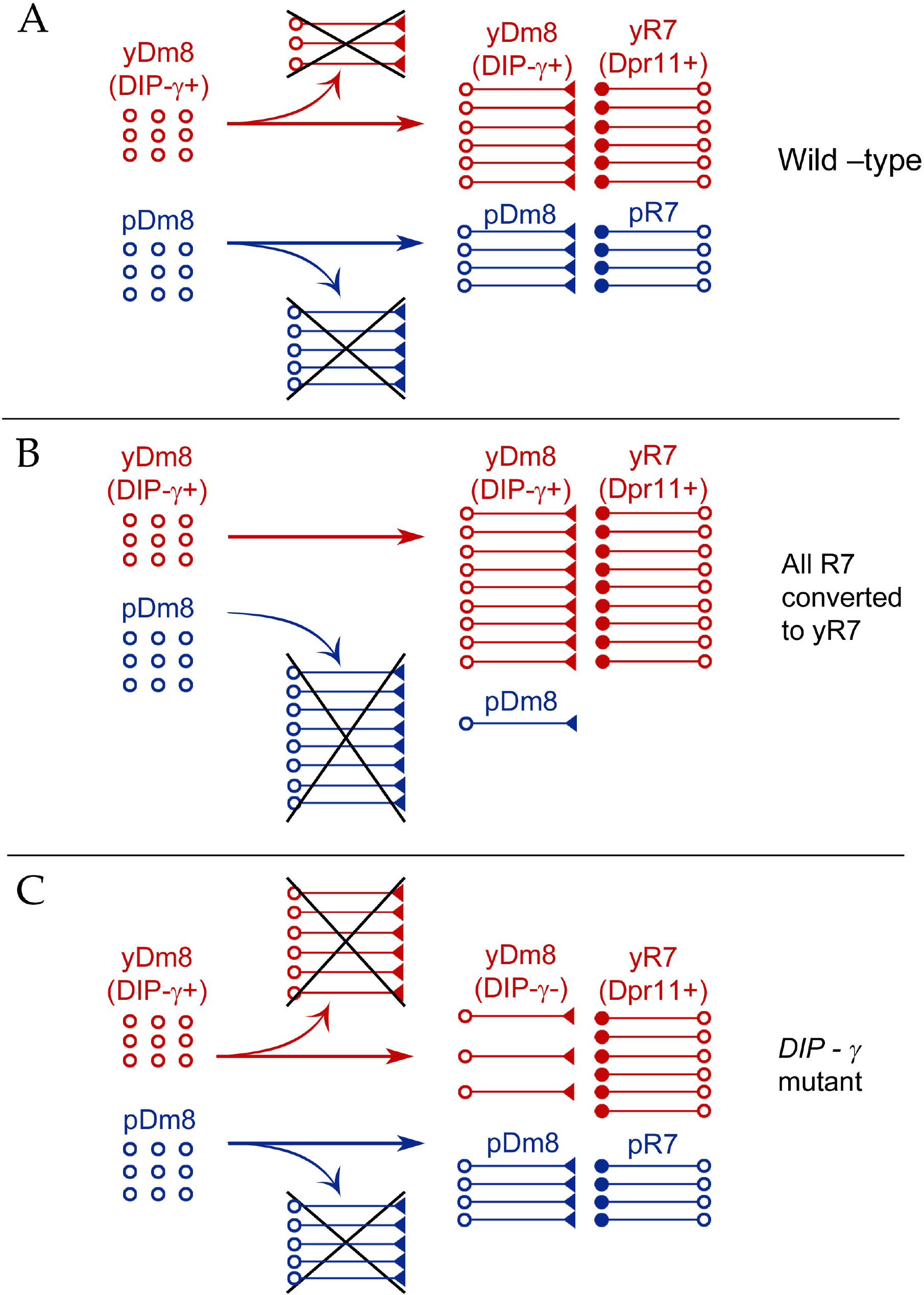
Model for Dm8 selection and survival. (A) In wild-type, yDm8 neurons (red) are produced in excess and are selected by yR7 PRs for connectivity and survival. pDm8 neurons (blue) are also shown as being generated in excess, and we have indicated their cell numbers prior to selection as being the same as yDm8, but this is arbitrary, since we do not know how many pDm8 are born. Unselected yDm8 and pDm8 neurons die in the absence of neurotrophic support (indicated by Xs). ∼30% of the yDm8 neurons present at 15h APF die in wild-type (Figure 7A). (B) When all R7 PRs are converted to yR7 (in *ss^GOF^*) or when they all express Dpr11, more yDm8 neurons survive and almost all pDm8 neurons die (Figure 8). (C) In *DIP-*γ mutants, yDm8 neurons are not selected by yR7 PRs and more of them die. ∼75% of the yDm8 neurons present at 15h APF die in *DIP-*γ mutants (Figure 7A). Similar results are observed for *dpr11* mutants. This means that some yR7 PRs remain uninnervated (Figure 6-figure supplement 2).

Dpr11 and DIP-γ are required for survival, because 50-60% of yDm8 die when either molecule is absent (Figures 6, 7A, 9). If all R7 are converted to yR7 (in *ss^GOF^*), or if they all express Dpr11, many more yDm8 neurons survive than in wild-type, showing that the numbers of Dpr11-expressing R7 are limiting for yDm8 survival in wild-type animals (Figures 8C, 9). This seems surprising, because a wild-type retina should contain 460-500 yR7 (Posnien et al., 2012), which is greater than the number of yDm8 we count in early pupae (Figure 7A). However, the actual yDm8 cell numbers may be larger due to incomplete penetrance of the markers, as discussed in Results.

yDm8 and pDm8 populations do not compete with each other in wild-type because they have different R7 partners. When the numbers of yDm8 neurons are increased by suppressing apoptotic cell death with DIAP in yDm8, pDm8 cell numbers do not decrease (Figure 6B). However, yDm8 and pDm8 neurons can be forced to compete by changing R7 subtype fate. When all R7 PRs are converted to yR7, or when they all express Dpr11, almost all pDm8 neurons die (Figures 8D, 9). Conversely, most yDm8 neurons die when all R7 PRs are converted to pR7 (in *dve^GOF^*)(Figure 8G).

pDm8 neurons are preferentially selected by pR7 PRs, using unknown molecular mechanisms, and pR7 PRs are needed to ensure pDm8 survival. This is demonstrated by the fact that pDm8 are unable to effectively fill vacant yR7 slots. For example, in *ss^GOF^, DIP-*γ animals, there are at least 180 yR7 slots that are made vacant by loss of yDm8, but the number of pDm8 neurons only increases by ∼30 (∼2-fold) in this genotype (Figures 8C-D).

### Selection of yDm8 neurons for survival occurs early, prior to synaptogenesis

Dpr11 is already expressed in a subset of R7 PRs at 15h APF, indicating that yR7 fate is determined by that time (Figure 7E). DIP-γγis expressed by 18h-20h APF in select Dm8 neurons apposed to Dpr11-expressing R7 PRs (Figure 7F; Figure 7-figure supplement 1A-B). At this time, R7 growth cones are positioned in the incipient layer (Kulkarni et al., 2016; Ting et al., 2005). This is consistent with earlier findings that single-cell Dm8 clones contact home column R7 growth cones by 17h APF (Ting et al., 2014). Taken together, these results indicate that yR7 and yDm8 have met and selected each other in the incipient layer by 15h-20h APF. Neurotrophic signaling mediated by Dpr11-DIP-γγinteractions is probably initiated around that time, because cell death of yDm8 in *DIP-*γγmutants begins to occur between 15 hr and 25 hr APF (Figure 7A).

There is widespread cell death in the medullary cortex as part of normal optic lobe development. This occurs in two phases, with the first phase spanning 0h-48h APF and peaking at 24h APF (Togane et al., 2012). In wild-type, ∼30% of the yDm8 neurons present at 15 h APF die by 45h APF, at which time yDm8 cell numbers are the same as those in adult. In *DIP-*γ mutants, ∼75% are lost by 45h APF. If apoptotic cell death is suppressed by DIAP, then the yDm8 population in adult remains the same as at 15h APF (Figure 7A). We suggest that the yDm8 neurons that die in wild-type are those that are in excess of the number that can be selected for survival by Dpr11-expressing yR7 PRs. Since most yDm8 neurons require DIP-γ for survival, we infer that the excess yDm8 neurons in wild-type die because they lack neurotrophic support mediated by Dpr11-DIP-γ interactions. However, we do not understand why yDm8 neurons begin to die earlier in *DIP-*γ mutants than in wild-type (Figure 7A). Perhaps DIP-γ not engaged by Dpr11 (which would be present in wild-type yDm8) allows cells to survive longer than when DIP-γ is absent. These selection events are independent of synaptogenesis between R7 and Dm8, which occurs only after 60h APF.

The DIP-α-expressing Dm4 neurons are also generated in excess, and their survival is regulated by interactions between DIP-α and its partners Dprs 6 and 10, which are expressed on presynaptic L3 neurons. In *DIP-α* or *dpr6 dpr10* mutants, about 50% of Dm4 and 20% of Dm12 neurons are lost. If cell death is blocked in wild-type, the cell numbers of Dm4 neurons in adults are increased by about 25% (Xu et al., 2018b). Thus, in these cases Dpr-DIP interactions play a similar role in suppressing cell death of postsynaptic neurons in the distal medulla. However, not all DIP-γ expressing neurons in the optic lobe require it for survival. A subset of lobula plate tangential cells (LPTCs) also express DIP-γ, and these do not die in *DIP-*γ mutants (unpublished data).

Our gain-of-function studies show that when Dpr11 is expressed in all R7, all pDm8 home columns are converted to yDm8 (Figures 8E-F, H), demonstrating that expression of Dpr11 in R7 PRs that are otherwise of the pale subtype is sufficient for selection of yDm8. However, we were unable to determine whether Dpr11 is also necessary for yDm8-yR7 pairing by analysis of LOF mutants. We did not observe mistargeting of surviving yDm8 in *dpr11* mutants to pR7 columns, most likely because pR7 slots are occupied by pDm8 neurons (Figure 8-figure supplement 2A-C). Another factor that could contribute to accurate targeting in *dpr11* mutants is that there may be a second mechanism for support (and possibly selection) of yDm8 by R7 that is independent of Dpr11-DIP-γγinteractions. This is suggested by the fact that ∼70% of yDm8 are lost in *sev* mutants, but only ∼50% in *DIP-*γγand *dpr11* mutants (Figures 8C, 6A).

### Effects of *DIP-*γ and *dpr11* mutations on yDm8 arbor morphology

A typical wild-type yDm8 arbor has a thick distal dendritic projection (sprig) that extends along the home column yR7 terminal, usually reaching M4. In *DIP-*γ and *dpr11* mutants, the distribution of sprig diameters for yDm8 neurons is shifted toward smaller values (Figure 4H), suggesting that mutant sprigs may have a reduced capacity to form synapses with yR7 in M4 and M5. Synapses that would have been in M4 and M5 might then be found in the base of the arbor in M6. EM reconstruction data for columns D and E supports this idea. R7 output synapses (T-bars) are mainly in M4 and M5 (on the sprig) in column E, which has a Dm8 with a thick sprig (Figures 2A, 3A, inset in 3C). In column D, whose Dm8 has a very thin sprig, R7 T-bars are located in M6 (in the main arbor) (Figures 3B, inset in 3D).

The effects of Dpr11 and DIP-γ on yDm8 sprig morphology may involve a different pathway from that involved in cell death. Although cell death would have occurred in the *DIP-*γ and *dpr11* mutant animals, these flipouts were analyzed in adults, in which the population has stabilized and the remaining yDm8 neurons are not in the process of dying. Sprig morphology defects might be a consequence of a loss of Dpr11-DIP-γ-mediated adhesion between yR7 and yDm8, since *dpr11* and *DIP-*γ mutants are similarly affected.

We had earlier reported an “overshoot” phenotype in *dpr11^GFP^/Df* mutants in which some R7 terminals had processes extending beyond M6 into proximal medulla layers(Carrillo et al., 2015). Overshoots were labeled by an overexpressed truncated form (Brp-short) of the active zone protein Bruchpilot (Brp)(Berger-Muller et al., 2013). However, this phenotype was not observed in the *dpr11^CRISPR^* null mutant when endogenous Brp puncta were labeled with the STaR method (Xu et al., 2018b). To determine whether this discrepancy was due to the *dpr11^GFP^/Df* genotype or to the Brp-short marker, we analyzed *dpr11^CRISPR^* homozygotes using Brp-short and observed a significant increase in yR7 overshoots as compared with the heterozygous control (Figure 8-figure supplement 2D). However, the interpretation of this Brp-short phenotype is unclear, as it does not correspond to a shift in the distribution of endogenous Brp puncta into layers proximal to M6.

### A circuit for wavelength discrimination defined by Dpr11 and DIP-γγexpression

Tm5a, Tm5b, and Tm5c are output neurons for R7 and R8 circuits. Their dendrites receive direct synaptic input from R7, R8, and Dm8, and their axons relay signals to the lobula (Gao et al., 2008; Karuppudurai et al., 2014; Takemura et al., 2013; Takemura et al., 2015). It has been hypothesized that Tm5a/b/c cells are analogous to vertebrate retinal ganglion cells, and that true color vision (intensity-independent hue discrimination) involves interpretation of Tm inputs by circuits in the lobula. Chi-Hon Lee and his colleagues developed a learning paradigm in which flies discriminate between equiluminant blue and green light, which are differentially sensed by y (Rh6) and p (Rh5) R8. Four classes of Tm neurons (Tm5 a/b/c and Tm20) make redundant contributions to this response (Melnattur et al., 2014).

The absorption spectra of Rh4 (yR7) and Rh3 (pR7) extensively overlap (Figure 3H), and it is unlikely that a learned color vision paradigm could be developed that would distinguish between Rh4 and Rh3 inputs. Nevertheless, the existence of discrete y and p R7 channels suggests that the fly utilizes them to make discriminations among UV wavelength inputs. To make such discriminations would require that the y and p channels have different synaptic circuits with distinguishable outputs. Our analysis of the patterns of synaptic connections defined by the EM reconstruction of the medulla (Takemura et al., 2013; Takemura et al., 2015) indicates that yR7 and yDm8 both preferentially synapse onto Tm5a, while pR7 and pDm8 synapse onto Tm5b (Table 1). Tm5a expresses DIP-γ, while Tm5b does not (Cosmanescu et al., 2018), so Dpr11-DIP-γ interactions might be involved in specifying both yR7-yDm8 and yR7-Tm5a connections. Four of the seven Dm8 neurons, and three of the eight R7 PRs, also make a few synapses onto Tm5c neurons, which are required for innate UV preference (Karuppudurai et al., 2014)(Table 1-table supplement). There is no specificity for y *vs*. p in the connections to Tm5c.

We visualized yR7-yDm8-Tm5a (yellow) and pR7-pDm8-Tm5b (pale) circuits using 3D renderings of cells from the EM reconstruction (Takemura et al., 2015)(Figures 3A-E and associated videos). The yDm8-E sprig and Tm5a-E dendritic branch are both wrapped tightly around the yR7-E terminal, and most yR7 T-bars are apposed to the sprig and branch within layer M5. Dm8 arbors contain both pre- and postsynaptic elements, and therefore both receive inputs and emit outputs. yDm8-E T-bars are distributed between the sprig and the main arbor in M6 (Figure 3C). Remarkably, our 3D renderings of wild-type yDm8 and yR7 neurons from ExM (Figure 5 and associated videos) are almost superimposable on yDm8-E and yR7-E from the EM reconstruction (Figure 5-figure supplement 1). This allows us to interpret yDm8 ExM phenotypes in *DIP-*γ mutants (Figure 5, Figure 5-figure supplement 2) by reference to the EM reconstruction.

R7 is histaminergic, and inhibits both Dm8 and Tm5a/b (Figures 3A-B), usually through polyadic synapses where a single R7 T-bar is apposed to both a Dm8 and a Tm5a/b postysnapse (Figures 3F-G). Dm8 is glutamatergic, while Tm5a and Tm5b are cholinergic (Davis et al., 2018; Gao et al., 2008; Karuppudurai et al., 2014). This suggests that yR7 input might cause inhibition of Tm5a in two ways: by direct inhibition and by inhibiting the glutamatergic input (probably excitatory) of yDm8 onto Tm5a. Conversely, pR7 input could preferentially inhibit Tm5b by both direct and indirect mechanisms. There might be timing differences between direct and indirect inhibition.

UV inputs will always stimulate both y and pR7 channels, because the Rh4 and Rh3 absorption maxima differ by only 20 nm. However, longer-wave inputs that activate Rh4 on yR7 more than Rh3 on pR7 might cause more inhibition of Tm5a than of Tm5b, while the reverse would be true of shorter-wave inputs that preferentially activate Rh3 on pR7 (Figure 3H). Combining direct (R7→Tm5a/b) and indirect (R7→Dm8→Tm5a/b) inhibition of Tm5a and Tm5b outputs might amplify these effects, depending on the relative timing of R7 and Dm8 inputs. This model suggests that neurons in the lobula or elsewhere that can read the ratio of Tm5a to Tm5b output mediate UV wavelength discrimination. Tm5a/b/c and Tm20 as a group synapse onto many lobula neuron types, but the specific partners of Tm5a and Tm5b are mostly unknown. Interestingly, however, all Tm5a but only half of Tm5b neurons synapse onto LT11 lobula projection neurons (Lin et al., 2016). Flies with silenced LT11 neurons have reduced phototaxis toward blue light (Otsuna et al., 2014).

In conclusion, our results show that Dpr11 and DIP-γ expression patterns define a yR7-yDm8-Tm5a circuit that should preferentially respond to longer-wavelength UV input. Neurotrophic signaling triggered by engagement of Dpr11 on yR7 by DIP-γ on yDm8 helps to build this circuit by ensuring that each yDm8 has a home column yR7.

## MATERIALS AND METHODS

### Drosophila genetics

Heterozygote controls were used in all experiments for determining cell numbers. *DIP-*γ *^GFP^* reporter (indicated in all graphs as *DIP-*γ ***^-^****^/+^*) is an insertion in the 5’ UTR intron and has reduced or no protein expression. Thus, this line serves as a mutant as well as a reporter of *DIP-*γ transcript.

For eye-specific transgenic RNAi, we screened 3 *dpr11* RNAi lines by crossing to a line which had one copy of *dpr11* removed to increase the effectiveness of the RNAi line: *lGMR-Gal4; dpr11^null,^ DIP-*γ *^MI03222-GFP^*. GD2343 (VDRC) had the strongest phenotype and was used in the paper.

### Generation of UAS-transgenic flies

cDNA encoding Dpr11 and DIP-γ were cloned from pOT2 GH22307 for Dpr11 and pOT2 GH08175 for DIP-γ into pUAST attB vector using standard molecular biology techniques. 5’ UTRs for both genes contained many upstream ATG codons; we made deletions of the 5’ UTRs so that the ATGs of the proteins were the first ATGs in the mRNAs, in order to increase expression of the transgenes (sequences available on request). Transgenes were injected into embryos (Rainbow transgenics). UAS-Dpr11 and UAS-DIP-γ both used the attP40 (2L) landing site.

### Antibodies

The primary antibodies used were as follows: anti-DIP-γ (guinea pig, 1:200) and anti-Dpr11 (rabbit, 1:150) were gifts from C. Desplan. Anti-Pros MR1A (mouse 1:4), anti-Elav 7E8A10 (rat, 1:10), anti-Dac 2-3 (mouse 1:50), anti-chaoptin 24B10 (mouse 1:20) were obtained from Developmental Studies Hybridoma Bank (University of Iowa, IA). Commercial antibodies were used as follows: Rabbit anti-RFP (Rockland, 1:500), rabbit anti-GFP (Thermo fisher Scientific, 1:500), chicken anti-GFP (Aves labs, 1:500), mouse anti-myc 9E10 (Abcam, 1:500) and chicken anti-beta-galactosidase (Abcam, 1:1000). Secondary antibodies were obtained from Thermo-fisher Scientific and used at 1:500.

### Immunohistochemistry

Eclosed flies (less than 3-days old) were dissected in phosphate buffer saline (PBS) and fixed in 4% paraformaldehyde in PBS with 0.2% Triton-X-100 (PBT) for 20 min at room temperature. Brains were washed overnight in PBT, followed by a two-day incubation at 4^0^C with primary antibody that was diluted in blocking buffer (5% normal goat serum in PBT). Samples were then incubated with secondary antibody (similarly diluted in blocking buffer) for two-days, followed by washing with PBT and PBS and stored in Vectashield.

For single cell flipouts, brains were given heat-shock at 50h APF for 10-20 min at 37^0^C and 1-day eclosed flies were dissected for staining as above.

For cell counts and column analyses, optic lobes were separated from the central brain and mounted top-down. Confocal images were acquired on Zeiss LSM800 or LSM700 microscopes with a 40xobjective. For cell counts and single-cell flipouts, optic lobes were imaged with 1.5µm and 0.8 µm z-sections, respectively. Single slices or maximum intensity projections were exported with Zen software (Zeiss) for image processing.

### Image processing

Images were processed with Adobe Photoshop.

### Quantitation and Statistics

Cell numbers were determined blind to genotype using Zen software to do manual counts. For all data (cell soma, columns and Brp overshoots), only 1 optic lobe per animal was used for counts and statistics. P-value was determined using Student’s unpaired t-test from Graphpad Prism. All data reported in graphs are mean +/-standard deviation.

yDm8 sprig parameters were measured using ImageJ software. To obtain measurement of the widest point of a sprig, a slice at which the image was most in focus was selected. The height was measured from the most distal point of the sprig to its most proximal point before the processes of the sprig connected back to the dendritic base. The Brp overshoots were analyzed blind using ImageJ software, by an individual unrelated to that experiment.

### ExM method

Single cell flipouts were processed for expansion microscopy after immunohistochemistry (and confirming the presence of flipouts) according to a protocol from Tim Mosca, based on (Mosca et al., 2017). Briefly, labeled brains were washed in 50% PBT-PBS mixture before proceeding with steps for anchoring proteins to the sodium acrylate matrix. Gelling chambers were made on coated slides with No.1 coverslips as bridge and brains were embedded in sodium acrylate gel. After incubation for 2 hours at 37^0^C, the samples were excised from the solidified gel and processed for digestion with proteinase K in 6-well plates. Gel fragments were expanded by replacing the digestion buffer with several washes of water. Finally, in preparation for confocal imaging, the gel fragments containing the expanded transparent brains were placed on a 24mm x 50mm No.1.5 coverslip that was coated with poly-l-lysine. Preps were always covered in water to make sure they did not dry out during imaging. Expanded brains were imaged on Zeiss LSM800 with 40x Objective 1.1 NA.

### 3D reconstruction of ExM samples

ExM images were processed for 3D reconstruction using Imaris software (Bitplane). Using surface rendering and fluorescence thresholding, we were able to obtain an accurate 3D structure of a yDm8 in wild-type and *DIP-*γ mutant. Each channel was recreated by adding an additional surface. The threshold for new surface detection was set as needed per image minimizing the addition of surface fragments not present in the original image. Each new surface was segmented in order to isolate the appropriate region of interest. Video recordings were captured at 10 frames per second for a total of 300 frames.

## Acknowledgments

We thank Maximilien Courgeon and Claude Desplan for sharing unpublished antibodies and fly lines and for discussions, Larry Zipursky for sharing unpublished fly lines, comments on the manuscript and discussions, and Tim Mosca for his ExM protocol. We thank Violana Nesterova for figure preparation, Shuwa Xu for comments on the manuscript and discussions during the course of the work and Namrata Bali for help with ExM. The *dpr11* RNAi stock was obtained from the Vienna Drosophila Resource Center (VDRC, www.vdrc.at). Stocks obtained from the Bloomington Drosophila Stock Center (NIHP40OD018537) were used in this study. Imaging was performed in the Biological Imaging Facility, with the support of the Caltech Beckman Institute and the Arnold and Mabel Beckman Foundation. Anti-Pros MR1A (mouse 1:4), anti-Elav 7E8A10 (rat, 1:10), anti-Dac 2-3 (mouse 1:50), anti-chaoptin 24B10 (mouse 1:20) were obtained from the Developmental Studies Hybridoma Bank (University of Iowa, IA). This work was supported by NIH grants RO1 EY028116 and R37 NS28182 to K. Z, and by the Howard Hughes Medical Institute (S-Y. T).<colcnt=3>

## Additional Information

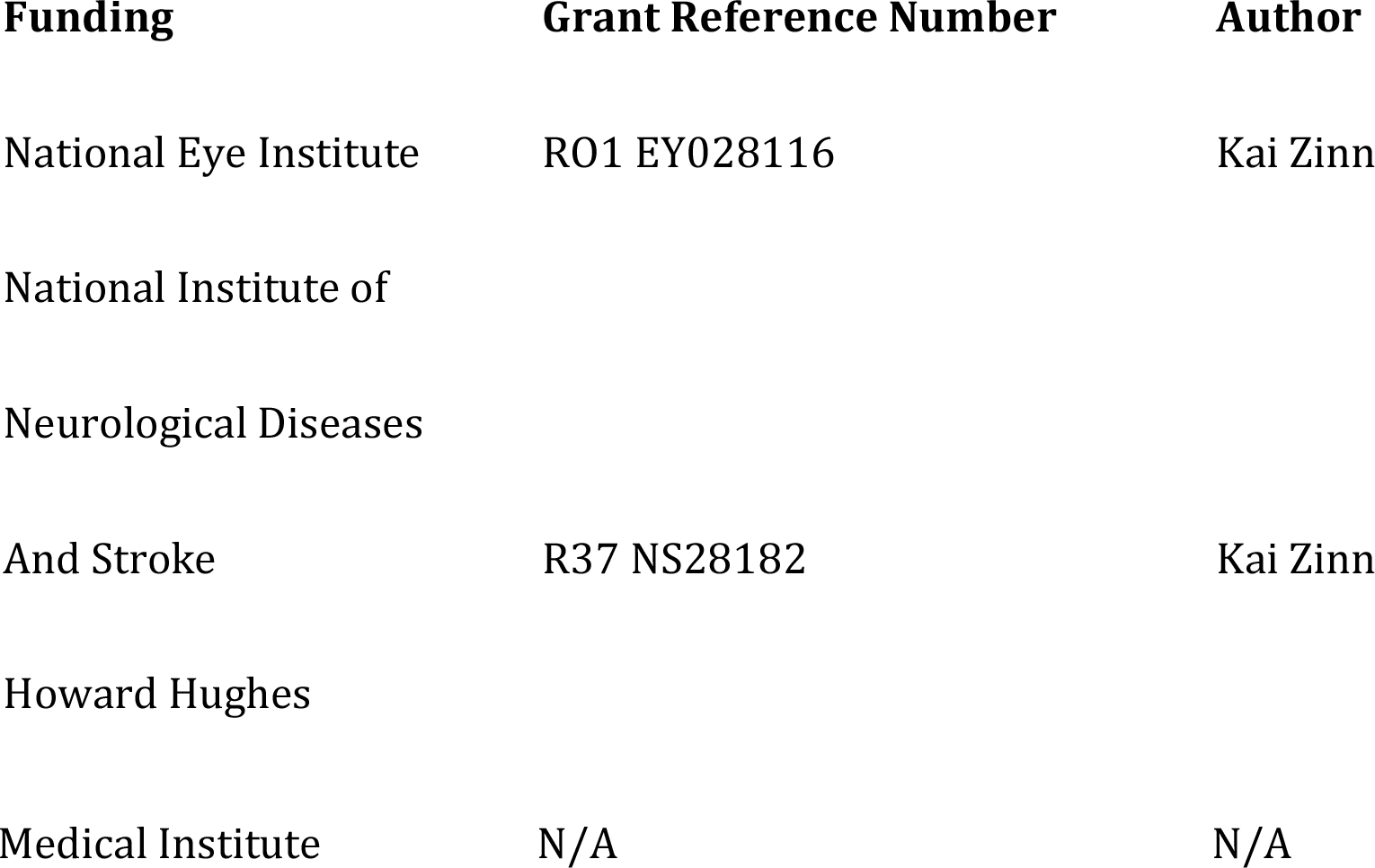

## Author contributions

Kaushiki P. Menon, Conceptualization, Supervision, Formal analysis, Validation, Investigation, Visualization, Methodology, Writing—original draft, Writing—review and editing; Vivek Kulkarni, Formal analysis, Validation, Investigation, Visualization, Methodology, Writing—review and editing; Shin-ya Takemura, Formal analysis, Validation, Investigation, Visualization, Writing—review and editing; Michael Anaya, Resources; Kai Zinn, Conceptualization, Supervision, Funding acquisition, Writing—original draft, Writing— review and editing, Project administration.

## VIDEOS

Figure 3 videos:

**1. Horizontal rotation of column E yellow circuit** Colors as in Figure 8

**2.** Vertical rotation of column E yellow circuit

**3. Horizontal rotation of column D pale circuit**

**4. Vertical rotation of column D pale circuit**

**5. Horizontal rotation of column B/home pale circuit**

**6. Vertical rotation of column B/home pale circuit**

## Figure 5 videos

**1.Horizontal rotation of an expanded yDm8 in wild-type**

Horizontal rotation of a yDm8 (cyan) with Chp labeled R7 PRs (magenta). A single yDm8 wild-type clone was expanded and surface rendered with Imaris software. Note that in addition to the major dendritic process (sprig) wrapping around the home column R7, two thinner dendritic processes extend distally along other R7.

**2. Vertical rotation of an expanded yDm8 in wild-type**

Vertical rotation of the same yDm8 clone.

**3. Horizontal rotation of an expanded yDm8 in *DIP-*γ mutant**

Horizontal rotation of a yDm8 (cyan) and a Chp labeled home column yR7 (magenta). A single yDm8 *DIP-*γ mutant clone was expanded and surface rendered with Imaris software. The thin process that is not in contact with the R7 is the yDm8 axon.

**4. Vertical rotation of an expanded yDm8 in *DIP-*γ mutant**

Vertical rotation of the same mutant yDm8 clone.

---

**Figure 2-figure supplement 1:**
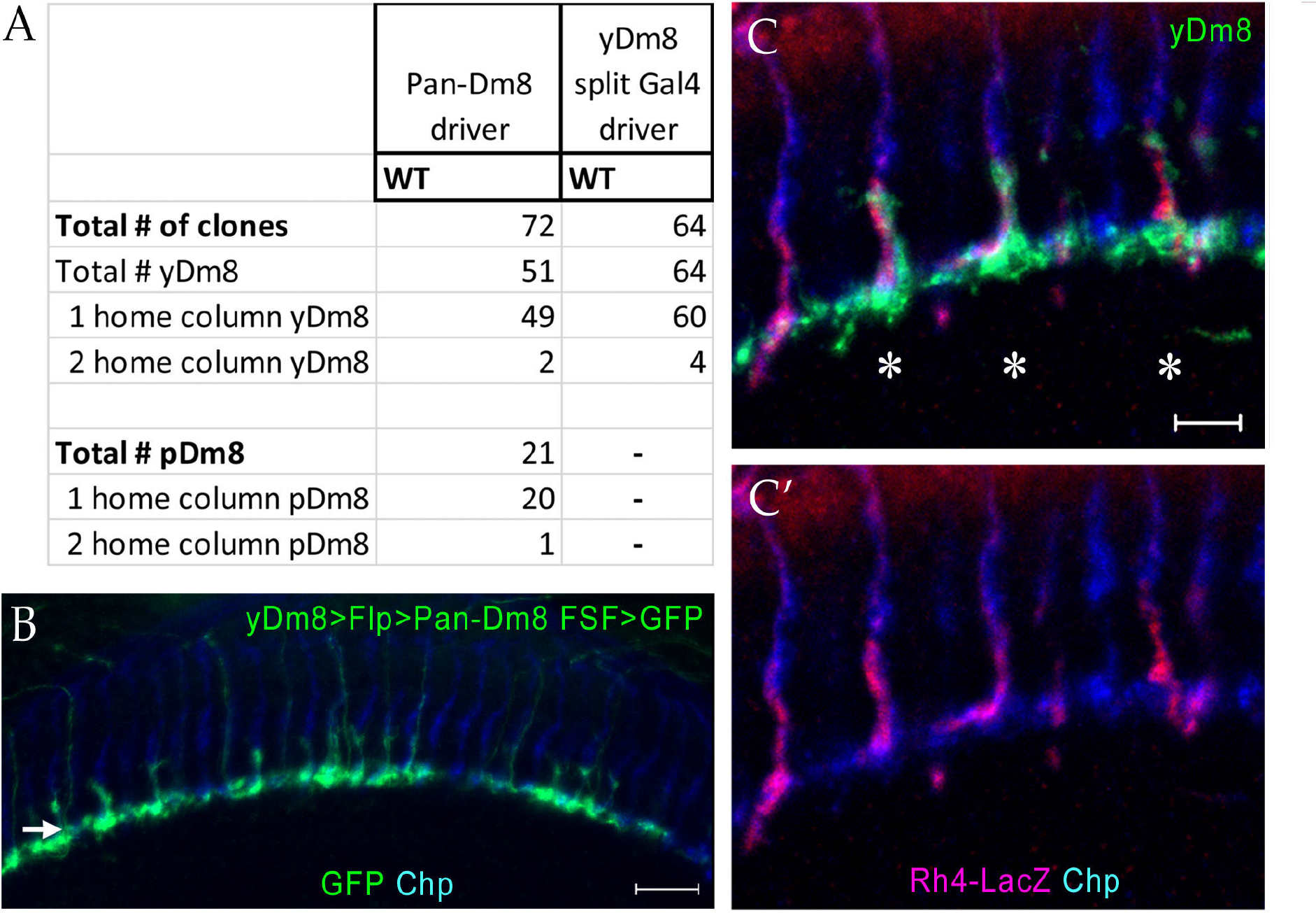
Quantitation of yDm8 and pDm8 flipout clones. (A) Quantitation of yDm8 and pDm8 flipout clones in wild-type: Pan-Dm8 driver: R24F06-Gal4 (Nern et al., 2015). yDm8 split-Gal4 driver: R24F06-p65.AD; DIP-γ-Gal4-DBD (This study). (B) yDm8 and pDm8 populations have independent origins. Merged panel of Figure 2G. The dendritic arbors of yDm8 neurons are labeled in flies carrying DIP-γ Gal4>Flp and pan-Dm8-LexA>LexAop-FRT-stop-FRT>GFP transgenes. pDm8 that are not labeled appear as gaps in the M6 layer (arrow), similar to those seen when yDm8 neurons only are labeled by *DIP-*γ*^GFP^* (Figure 2H’). Adult optic lobes were labeled with anti-GFP (green) and anti-Chp (blue). Maximum intensity projection; scale bar 10 µm. (C-C’) Horizontal view of a dense flipout showing 3 yDm8 clones (asterisks) with yR7 home columns labeled with Rh4-LacZ (red) and Chp (blue). Note the absence of sprig labeling on a pR7 column labeled with Chp only (blue). Maximum intensity projection; scale bar 5 µm.

**Figure 3-figure supplement 1:**
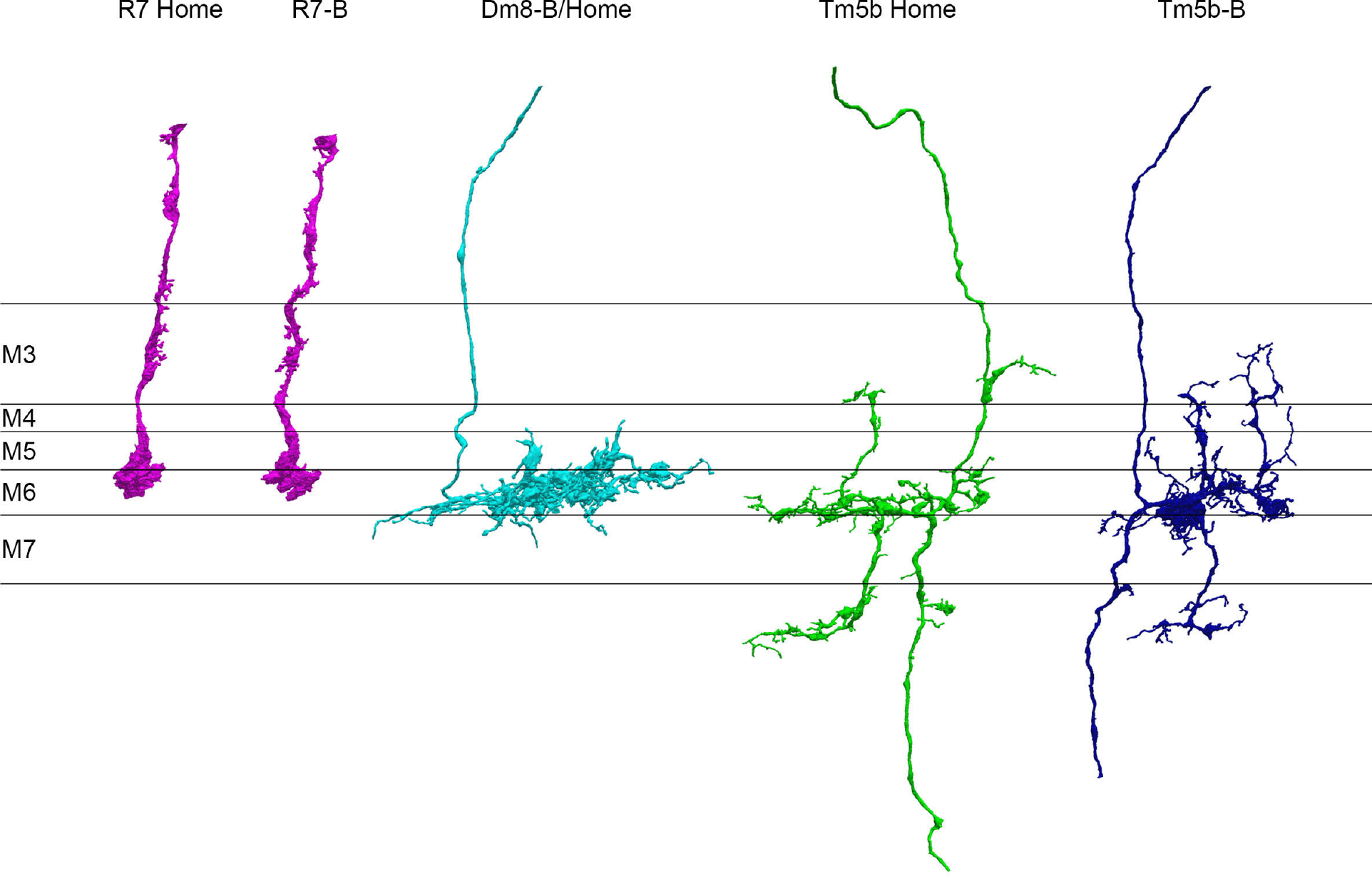
Separated cell profiles for the column B/home two-home column circuit.

**Figure 4-figure supplement 1:**
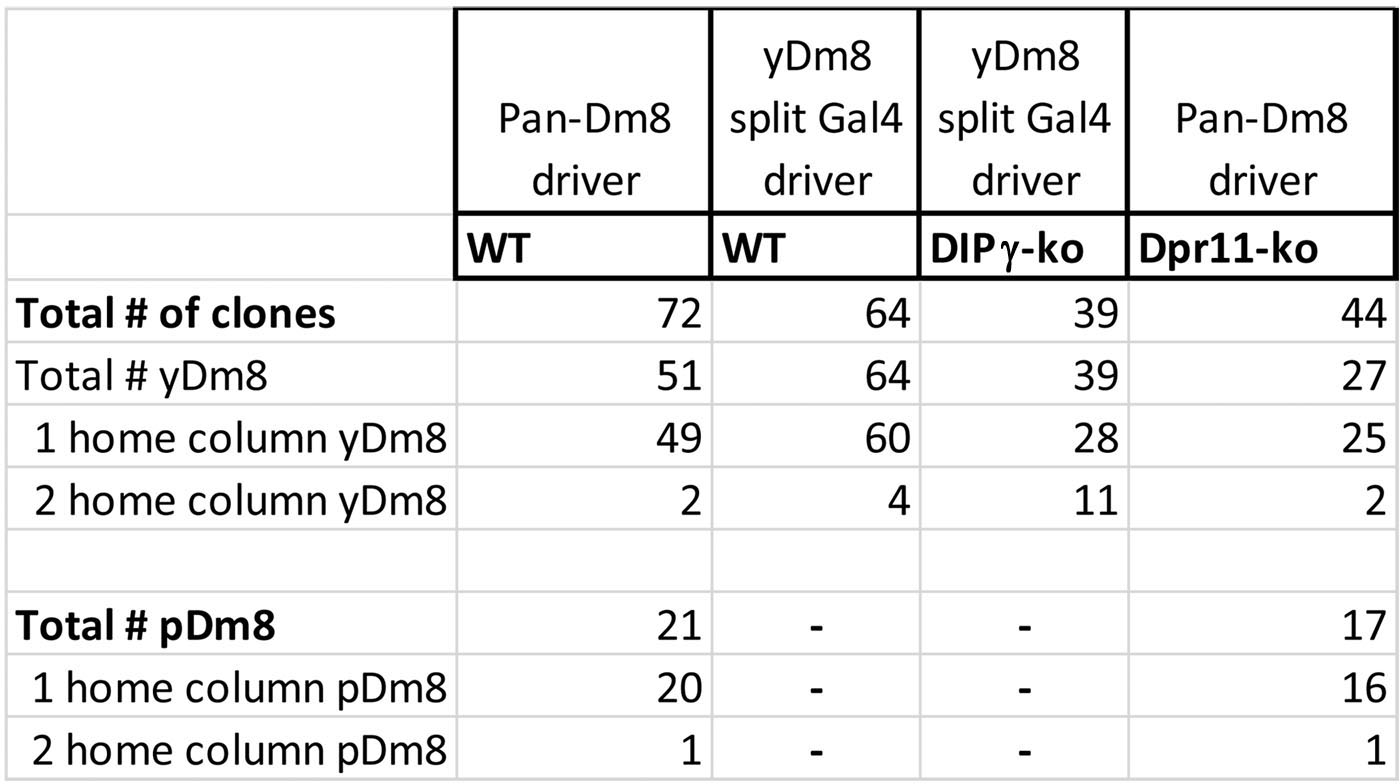
Frequency of two-home column yDm8 clones is increased in *DIP-*γ mutants. (A): The percentage of two-home column yDm8 clones in *DIP-*γ *^-/-^* mutants is 4.5-fold higher than in wild-type. Quantitation of flipout clones in wild-type, *DIP-*γ and *dpr11* mutants shown. Flipouts were generated with either pan-Dm8 driver *R24F06-Gal4* or the yDm8 split-Gal4 driver *R24F06-p65.AD; DIP-*γ*-Gal4-DBD*.

**Figure 5-figure supplement 1:**
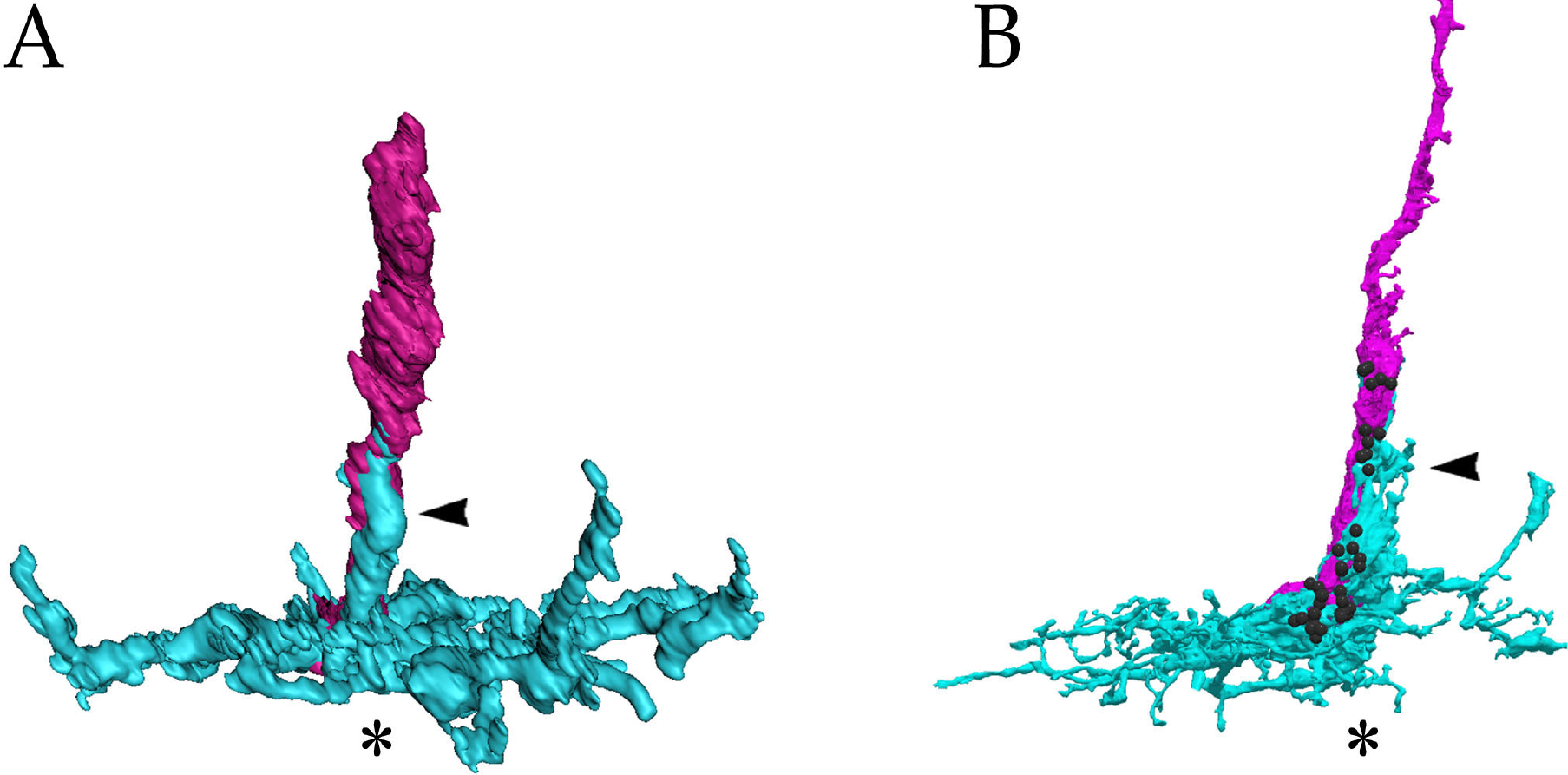
A comparison of the ExM rendering of the wild-type yDm8 and yR7 shown in Figure 5B, and the EM reconstruction of yDm8-E and yR7-E shown in Figure 2A. yDm8, cyan; yR7, magenta. Black balls indicate yR7 T-bars. Based on this comparison, we can infer that there are likely to be many R7-Dm8 synapses on the sprig in (A).

**Figure 5-figure supplement 2:**
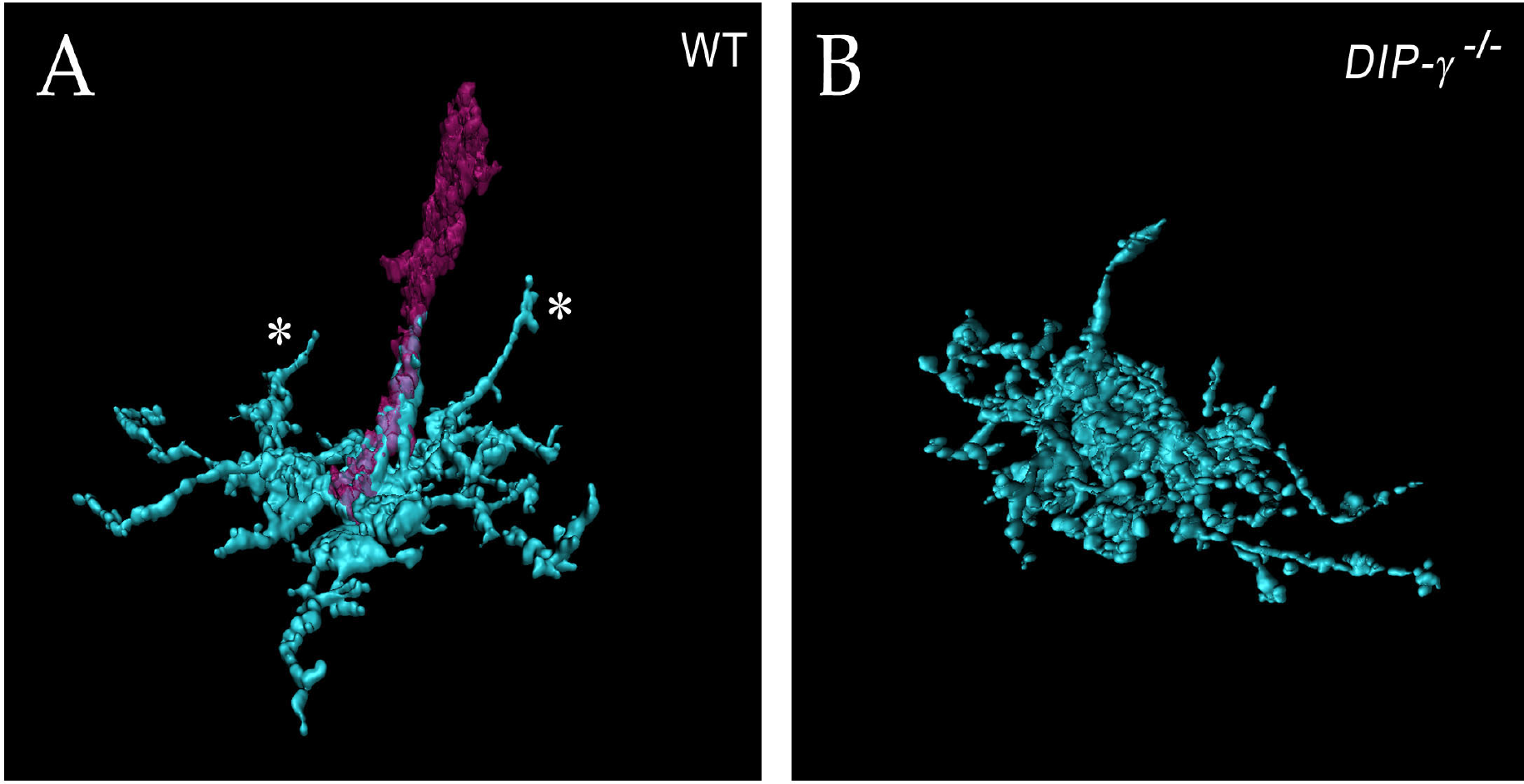
Other views of expanded yDm8 in (A) wild-type and (B) *DIP-*γ *^-/-^*. (A) The wild-type rendering is the same column as in Figure 5C, but with only the home column yR7 included. This allows clearer visualization of the other two dendritic projections (asterisks). (B) The mutant rendering is without the R7, allowing clearer visualization of sprig morphology.

**Figure 6-figure supplement 1:**
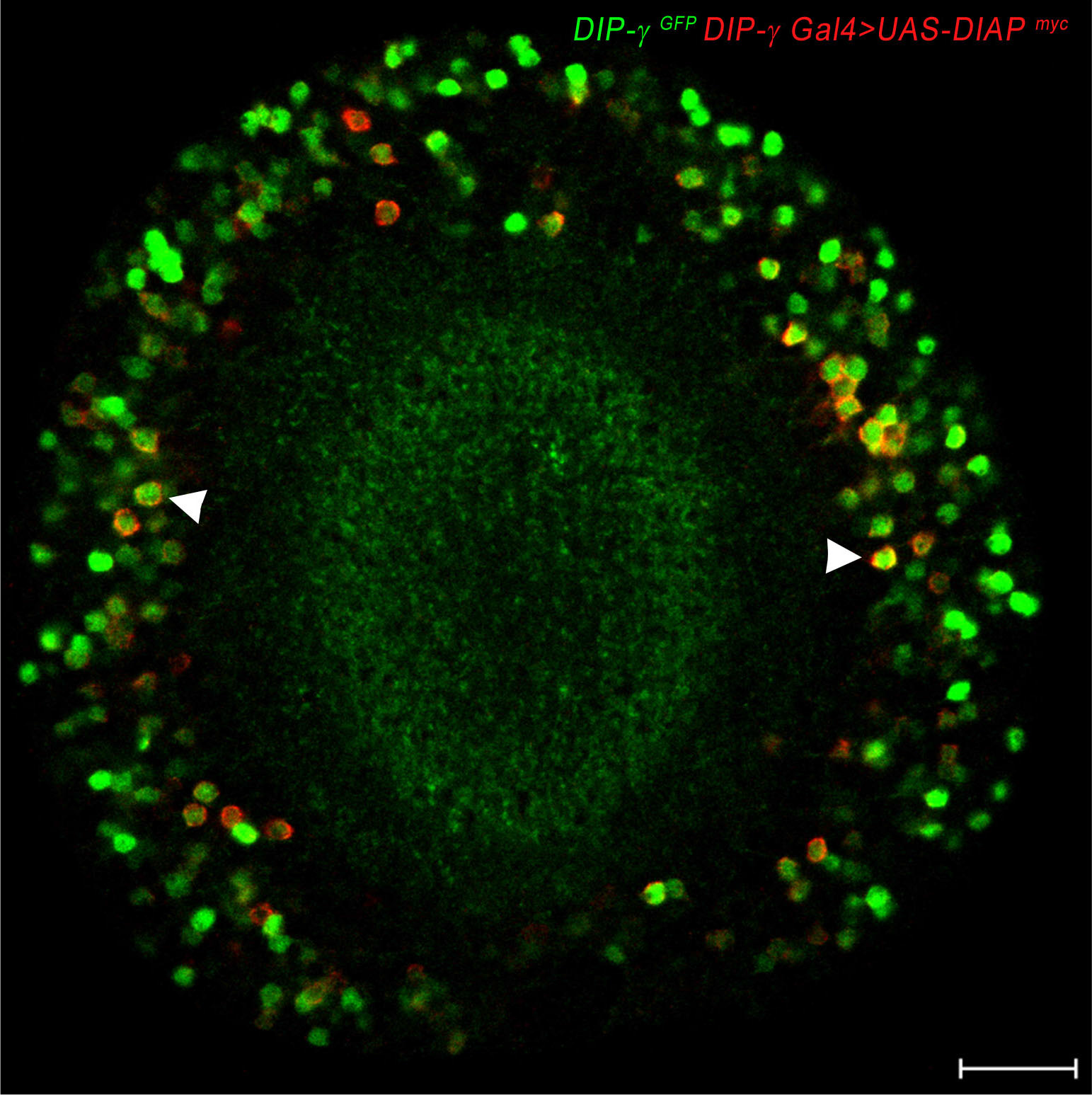
DIAP localizes to DIP-γ expressing cells. DIAP1 tagged with myc (red) driven with *DIP-*γ *Gal4* shows co-localization of DIAP1 and *DIP-*γ *^GFP^* reporter (green). Adult optic lobe labeled with anti-myc (red) and anti-GFP (green). Single section. Scale bar 20 µm.

**Figure 7-figure supplement 1:**
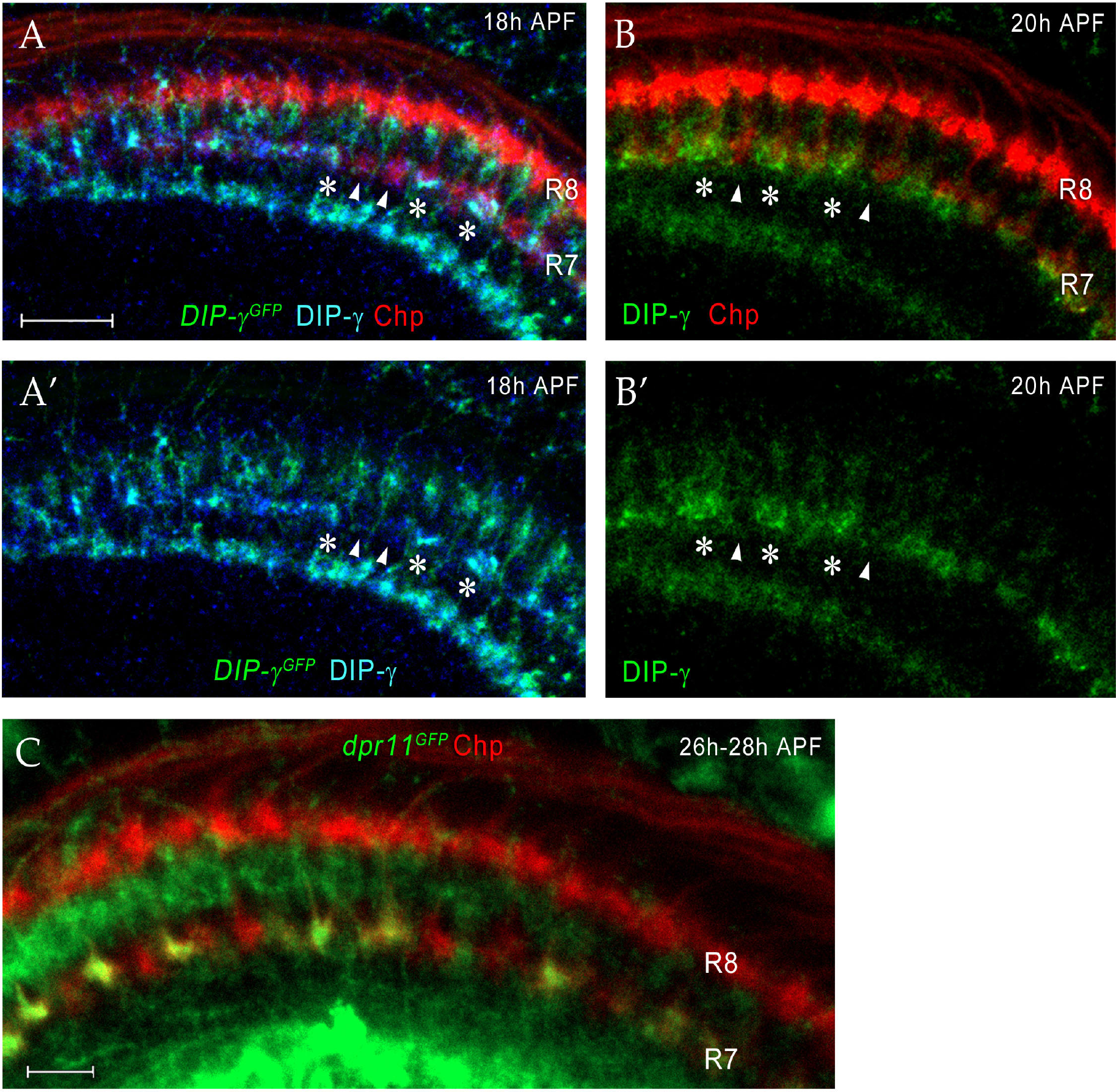
Dpr11 and DIP-γ are expressed in yR7 and yDm8, respectively, around the time yR7 selects yDm8 for survival. (A-A’) DIP-γ is expressed in yDm8 processes overlapping select R7 terminals by 18h APF. *DIP-*γ*^GFP^* reporter and DIP-γ antibody show the same pattern in the neuropil. Three R7 PRs with anti-GFP and anti-DIP-γ labeling are indicated by asterisks. There are gaps in DIP-γ labeling in the R7 incipient layer (one such gap covering two R7 terminals indicated by two arrowheads). *DIP-*γ*^GFP^* reporter labeled with anti-GFP (green), anti-DIP-γ (blue) and anti-Chp (red). (A’) is without the Chp labeling, for clearer visualization of the gaps. (B-B’) (B) is the same merged image seen in Figure 7F. (B’) shows the DIP-γ channel only for this image, and reveals gaps in DIP-γ labeling, implying that pDm8 are apposed to pR7 within the gaps (arrowheads). DIP-γ labeled yDm8 processes indicated by asterisks. Wild-type labeled at 20h APF with anti-DIP-γ (green) and anti-Chp (red). (C) *dpr11^GFP^* expression at 26h-28h APF labels yR7 but not pR7. *dpr11^GFP^* reporter labeled with anti-Chp (red) and anti-GFP (green). R7 terminals unlabeled by GFP are pR7 (red).

**Figure 8-figure supplement 1:**
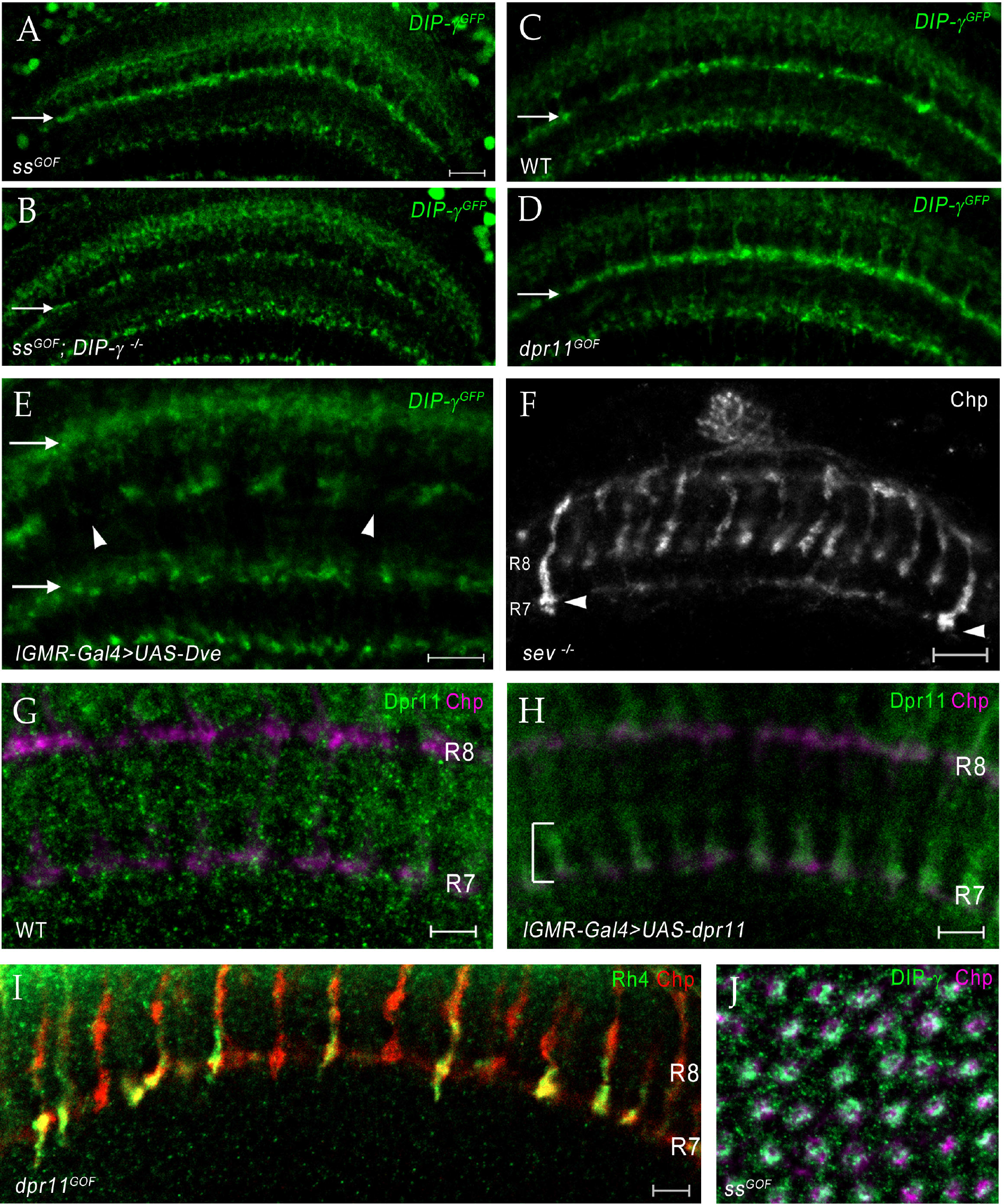
Changing R7 fate or expressing Dpr11 affects yDm8 and pDm8 survival. (A-D) Conversion of all R7 PRs to yR7 fate in *ss^GOF^* results in loss of pDm8 arbors. (A) *ss^GOF^; DIP-*γ *^-/+^* (B) *ss^GOF^*; *DIP-*γ *^-/-^* (C) Wild-type control (D*) lGMR>UAS-dpr11* (*dpr11^GOF^*)*. DIP-*γ *^GFP^* reporter in M6 layer (arrow) is shown for all panels. Gaps representing pDm8 arbors are present in wild-type control (C) but absent in *ss^GOF^* (A), where pR7 PRs have been replaced by yR7 PRs. Ectopic expression of Dpr11 in all PRs mimics *ss^GOF^* (compare panels A and D). (E) yDm8 arbors are lost when yR7 are converted to pR7 by expressing Dve in all PR. There are large gaps (arrowheads) in yDm8 labeling in layer M6 of the neuropil indicating extensive yDm8 cell death (compare to control (C), which has only small gaps). yDm8 arbors in M6 were examined with *DIP-*γ*^GFP^* reporter in *lGMR-Gal4>UAS-Dve*. The other layers in the distal medulla (M3) and in the proximal medulla that label with the reporter are unaffected in *dve^GOF^* (arrows in (E); compare to those layers in (C)). (A)-(E) single confocal slices; all scale bars are 10 µm. (F) Some R7 PRs remain in a *sev* “null” mutant. A single confocal slice of a *sev^14^/sev^14^* (putative amorphic mutant; see Flybase) adult, labeled with anti-Chp. There are two R7 axons that project to M6 visible in this slice (arrows). Scale bar, 10 µm. (G-H) Overexpressed Dpr11 can be detected on R7 terminals in *lGMR-Gal4>UAS-dpr11*. (G) *lGMR-Gal4* control (H) *lGMR-Gal4>UAS-dpr11*. Pupal optic lobes (∼43h APF) were labeled with anti-Dpr11 (green) and anti-Chp (magenta). Note distinct Dpr11 labeling (bracket) of R7 axons/terminals in (H), and its absence in (G). This antibody is weak and shows no labeling of specific neurons in wild-type. Maximum intensity projection; scale bar 5 µm. (I) Expression of Dpr11 in pR7 PRs does not convert them to the y fate. R7 terminals in *dpr11^GOF^* adult labeled with Rh4-lacZ and Chp. Note that 4 of the R7 terminals are labeled only by Chp and are therefore pR7. If Dpr11 expression in all PRs produced the same effect as *ss* expression, all R7 terminals would express both Rh4-lacZ and Chp, because they would all be y. Scale bar, 5 µm. (J) Ss overexpression (*ss^GOF^*) converts all R7 PRs to yR7, and converts almost all columns to yellow due to selection of yDm8, which ensures their survival. Cross-section view of medulla neuropil at 54h APF labeled with anti-Chp (magenta) and anti-DIP-γ (green). Compare to Figure 8F. Maximum intensity projection.

**Figure 8-figure supplement 2:**
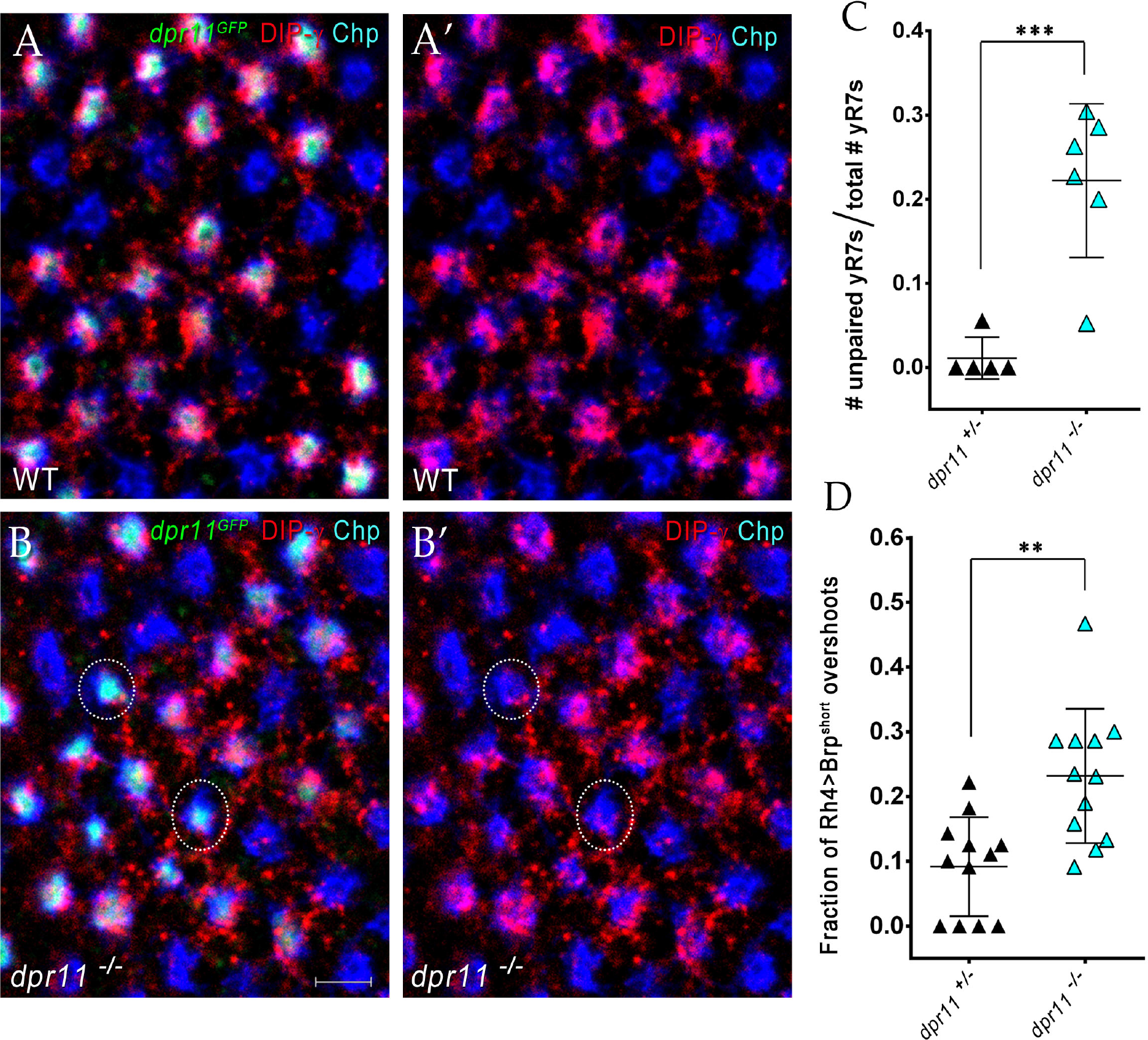
yDm8 arbors do not innervate pR7 home columns in *dpr11* mutants. (A)-(B): Surviving yDm8 do not mistarget to pR7 home columns in a *dpr11* mutant. Mid-pupal optic lobes of *dpr11^GFP^* heterozygote reporter line (WT) and *dpr11^GFP/-^* (mutant) were labeled with anti-GFP for yR7 (green), anti-DIP-γ for yDm8 (red) and Chp for all PRs (blue). (A) and (B) show all three channels, and (A’) and (B’) show only red and blue. A yR7 column labeled by DIP-γ is defined as one in which there are red pixels directly on top of the blue and green Chp and *dpr11^GFP^* labeling. Note that all yR7 columns with green labeling in (A) (these appear white) are labeled by both red and blue in (A’). However, in *dpr11* mutants, some yR7 columns (circled in (B) and (B’)) with green labeling have no red DIP-γ labeling on top of the column, indicating that these are vacant yellow columns that have no yDm8. These are quantitated in (C). All *dpr11* mutant columns are either blue (pR7-Chp only), or blue+red+green (Chp+DIP-γ+*dpr11^GFP^*), showing that no yDm8s mistarget to pR7s (*i.e*., there are no red+blue columns in (B)). (C): Some yR7 home columns are uninnervated in *dpr11* mutants due to yDm8 cell death. Pupal optic lobes of *dpr11^GFP^* reporter line (*dpr11^+/-^*) and *dpr11^-/-^* were labeled with anti-GFP for yR7, anti-DIP-γ for yDm8 and Chp for all PRs. Number of yR7 columns in the medulla were quantitated in 6x6 grids drawn on images of cross-section views. Number of yR7 columns without yDm8 partners was determined by counting how many *dpr11^GFP^* labeled yR7 did not have yDm8 labeling adjacent to them. Graph shows mean +/-std. deviation and unpaired Student’s t-test p-values. In these grids, we observed no mispairing of yDm8 with pR7, which would be demonstrated by finding anti-DIP-γ labeling adjacent to R7 columns that were unlabeled by anti-GFP. *dpr11^-/+^* 0.011+/-0.02, *dpr11^-/-^* 0.22+/-0.09, ***p=0.0008 (D): yR7 overshoots detected with a truncated Brp marker are increased in the *dpr11* null mutant. We repeated our previously published analysis of yR7 overshoots in *dpr11* mutants using the CRISPR-generated *dpr11* null allele instead of *dpr11^GFP^/Df* (Carrillo et al., 2015; Xu et al., 2018b). We used the same reporter as before, Rh4 driving a truncated version of Brp (Brp-short; (Berger-Muller et al., 2013)) and determined the number of overshoots in which Brp-short labeling was observed beyond (proximal to) M6. The quantitation was done blind by a person not involved in the experiment. Graph shows mean +/-std. deviation and unpaired Student’s t-test p-values. *dpr11^-/+^* 0.09+/-0.08, *dpr11^-/-^* 0.23+/-0.1, **p=0.0011

**Table 1-table supplement.**
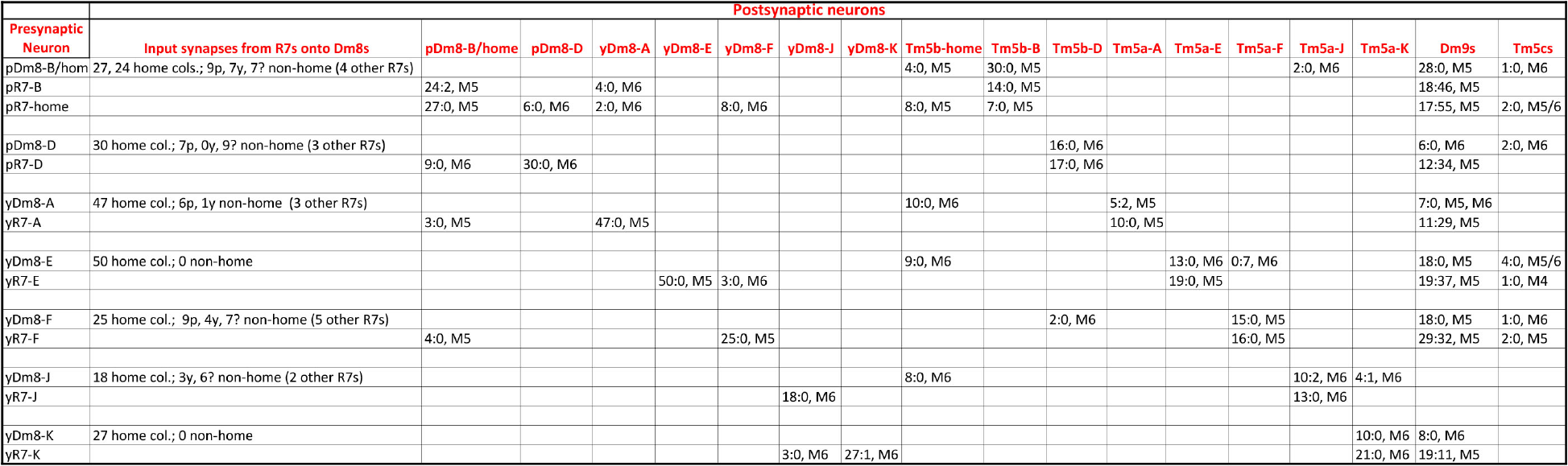
A table listing non-home column R7 synapses, and R7/Dm8 synapses with Tm5c and Dm9, in addition to the information in Table 1. Column 2 indicates the identities of the input synapses from R7 onto each Dm8. The reconstructed volume is unlikely to cover the entire arbor of these Dm8 neurons, since each Dm8 can contact 13-16 R7 (Gao et al., 2008). However, column 2 shows that within the reconstructed columns there is no apparent specificity in synaptic connectivity between y and p R7 and Dm8 neurons outside of their home columns. For example, pDm8-B/home receives synapses from 4 non-home column R7, at least 2 of which are yR7, while yDm8-A receives synapses from 2 non-home column pR7 and a yR7. Even the Dm8 at the center of the reconstruction (Dm8-B/home) only receives synapses from 4 non-home column R7 PRs, so many Dm8 contacts with R7 columns may not be associated with input synapses from R7. Also note that (Karuppudurai et al., 2014) found that Tm5a dendrites specifically associate with yR7 axons, but they stated that Tm5b dendrites have no specificity for association with y vs. p R7 axons. Our analysis of the EM reconstruction, however, shows that all 3 pR7 PRs synapse only onto Tm5b and not Tm5a. The conclusion of (Karuppudurai et al., 2014) likely arises from the fact that Tm5b usually has 2 distal dendritic projections. One of these always arborizes with a pR7 axon and receives the pR7 input. The other one can arborize with either a y or a p R7, but does not receive yR7 input, at least not for this set of columns. Four of the 7 Dm8, and 3 of the 8 R7 PRs, also have synapses onto Tm5c neurons. There is no specificity for y *vs*. p in the connections to Tm5c. Both R7 classes have input and output synapses with Dm9, and both classes of Dm8 have output synapses onto Dm9.

